# Ecological speciation within the *Phytophthora* genus

**DOI:** 10.1101/096842

**Authors:** M. F. Mideros, D. A. Turissini, N. Guayazán, G. Danies, M. Cárdenas, K. Myers, J. Tabima, E. M. Goss, A. Bernal, L. E. Lagos, A. Grajales, L. N. Gonzalez, D. E. L. Cooke, W. E. Fry, N. Grünwald, D. R. Matute, S. Restrepo

## Abstract

Over the past few years, symptoms akin to late blight disease have been reported on a variety of crop plants in South America. Despite the economic importance of these crops, the causal agents of the diseases belonging to the genus *Phytophthora* have not been completely characterized. In this study, we used an integrative approach that leveraged morphological, ecological, and genetic approaches to explore cryptic speciation within *P. infestans sensu lato*. We described a new *Phytophthora* species collected in Colombia from tree tomato (*Solanum betaceum*), a semi-domesticated fruit. All morphological traits and population genetic analyses, using microsatellite data and a reduced representation of single nucleotide polymorphism (SNP) data, support the description of the new species, *Phytophthora betacei* sp. nov. We have demonstrated that ecological differences are important in the persistence of *P. infestans* and *P. betacei* as genetically isolated units across an overlapping area in the northern Andes.

## Introduction

Oomycetes represent an opportunity to understand microbial speciation (Restrepo *et al.* 2014). Their ecological characteristics have been extensively studied to reveal a wide diversity of ecological niches (e.g Soanes *et al.* 2007). Particular emphasis has been placed on the study of plant pathogens for which ecological speciation seems to be a common process due to specialization to particular host species (Harrington *et al.* 2002; Tellier *et al.* 2010). Furthermore, oomycetes show an unprecedented plasticity in terms of genome size and ploidy (Haas *et al.* 2009a; Yoshida *et al.* 2013), which could influence rates of speciation and extinction (Santini *et al.* 2009; Wood *et al.* 2009; Muir & Hahn 2015; Puttick *et al.* 2015). Despite the hypothesized species richness of the group (Restrepo *et al.* 2014), no analytical framework currently delimits species boundaries in oomycetes.

Molecular taxonomy can use DNA sequences to identify and delimitate species that are not amenable to genetic crosses (Roe *et al.* 2010). The premise of these approaches is to identify discrete genetic groups that have ceased genetic exchanges with other groups. The number of studies defining species in this way has increased recently, mainly due to the ease of obtaining information on population-level DNA variation (Roe *et al.* 2010; Singh *et al.* 2015). However, this approach has inherent limitations. Gene genealogies tend to overestimate the number of species as the population structure within a species may be mistaken for species boundaries (Dettman *et al.* 2003). Furthermore, multi-locus species delimitation, relying on reciprocal monophyly and strict genealogical congruence, may fail to differentiate among recently diverged lineages (Hickerson *et al.* 2006; Knowles & Carstens 2007; Shaffer & Thomson 2007).

The most notable genus within oomycetes, *Phytophthora*, includes pathogens that infect a broad range of hosts in both agricultural and natural environments, causing adverse economic consequences (Erwin & Ribeiro 1996; Duncan 1999; Fry 2008; Forbes *et al.* 2013). To date, the genus *Phytophthora* comprises more than 150 recognized species, classified into 10 phylogenetic clades that are also supported by morphological and physiological characteristics (Blair *et al.* 2008; Kroon *et al.* 2012; Martin *et al.* 2014). Over the past few decades, the numbers of recognized species within most divisions of the genus *Phytophthora* have nearly doubled (Cooke *et al.* 2000; Kroon *et al.* 2012; Martin *et al.* 2012; Forbes *et al.* 2013). However, defining clear and objective species boundaries, as is the case for most oomycetes, remain a challenge in all *Phytophthora* clades.

Within the genus, *P. infestans* has become a “model system” because of its undoubted economic impact. This pathogen affects important crops, such as potato *(Solanum tuberosum)* and tomato *(Solanum lycopersicum)* (Haverkort *et al.* 2008; Visser *et al.* 2009), making it one of the most threatening plant disease agents in the world. Although *P. infestans* was once considered a single species (henceforth referred to as *P. infestans sensu lato*), it has been shown to be a species complex (Forbes *et al.* 2013). Four other species related to *P. infestans sensu stricto* have been identified over the last 35 years. *Phytophthora mirabilis* has been found in Central America (Galindo & Hohl 1985) infecting only *Mirabilis jalapa*, an ornamental and medicinal plant in the region. *Phytophthora ipomoeae* (Flier *et al.* 2002) infects two morning glory species endemic to the highlands of central Mexico, *Ipomoea longipedunculata* and *I. purpurea* (Flier *et al.* 2002; Badillo-Ponce *et al.* 2004). *Phytophthora phaseoli*, initially classified as *P. infestans* (Thaxter 1889), is distributed globally but infects only lima beans (*Phaseolus lunatus*). Due to host-preference studies, *P. phaseoli* was described as a different species and for over 60 years thought to be the closest relative of *P. infestans*. Genetic comparisons have also revealed the existence of a separate group composed of strains from Ecuador and Peru that are collectively called *P. andina* (Oliva *et al.* 2010). Phylogenetic hypothesis with both nuclear and mitochondrial markers, reveal this species as polyphyletic, suggesting that it might be a species complex (Adler *et al.* 2002, 2006; Kroon *et al.* 2004; Oliva ei *al.* 2010; Cárdenas *et al.* 2012; Forbes ei *al.* 2013, 2016; Goss *et al.* 2014; Lassiter *et al.* 2015). To date, *P. andina* is composed of the nuclear lineages EC-2 with two mitochondrial lineages, Ia and Ic; and EC3 (with one mitochondrial haplotype, Ia) (Oliva *et al.* 2010). A clear definition of the *P. andina* species is needed.

In this study, we used an integrative approach that leveraged morphological, ecological, and genetic approaches to explore cryptic speciation within *P. infestans sensu lato.* We found and defined a new species of *Phytophthora* infecting tree tomato (*Solanum betaceum*) in southern Colombia. Furthermore, we investigated the importance of host specificity in maintaining species boundaries within the *P. infestans sensu lato* species complex. Finally, we formally describe this new species as *Phytophthora betacei* sp. nov. For convenience, we refer to the species by using our proposed name for it throughout this article.

## Materials and methods

### Disease occurrence and collection of isolates

All *P. betacei* samples were collected in southern Colombia between 2008 and 2009 (Figure S1). We sampled three to four leaves from 10 randomly selected tree tomato plants per plantation that showed symptoms akin to late blight. In total, we sampled 34 locations from two Colombian states, Nariño and Putumayo (Figure S1). The initial collections comprised over 970 putatively infected leaves. One to three lesions per leaf were excised (~ 0.5 to 1 cm^2^) from the margin between necrotic and healthy tissues. Excised leaf pieces were surface-sterilized by submerging them in 70% ethanol for 20 to 30 s and then washed with sterile distilled water to remove excess ethanol (~ 10 sec). The leaf pieces were dried on a sterile paper towel and subsequently transferred to a selective medium prepared with tree tomato fruit (0.25 g of CaCO_3_, 0.5 g of yeast extract, 25 g of sucrose, 15 g of agar, and 100 ml of tree tomato extract, composed of 550 g of tree tomato fruit per liter of water). Subsequently, single zoospores were isolated from sporangia washed from the tree tomato medium with sterile distilled water. The sporangial suspension was adjusted to 2.0 × 10^3^ sporangia per ml, using a haemocytometer, and maintained at 4°C for 4 h to induce zoospore release before spreading 10 μl of the suspension onto 100-mm petri plates containing 10 ml of tree tomato medium. To better visualize zoospore germination, the medium was centrifuged at 8,000 rpm, and only the supernatant was used. The plates were incubated at 18°C for approximately 24 h before individual zoospores, observed through a stereoscope, were picked using a sterile syringe and placed onto fresh tree tomato medium.

In total, we successfully isolated 128 *P. betacei* strains. All single zoospore isolates were cultured in a tree tomato medium for 7 to 15 days at 18°C and then stored in the *Phytophthora* collection of the Museum of Natural History at Universidad de los Andes. Isolate P8084 was selected as the *P. betacei* holotype. This strain is maintained in culture, as well as cryopreserved with 1% glycerol, and deposited in the Museum of Natural History under accession number Andes-F 1172. All the other *P. betacei* isolates collected in this study are deposited in the same museum under accession numbers Andes-F 1081 to Andes-F 1207 (Table S1).

### Phylogenetic analyses and population genetics

#### DNA extraction and sequencing

The mycelia of each *Phytophthora* strain were grown in a liquid Plich medium (Van der Lee T *et al.* 1997) for 15 days at 20°C. Subsequently, mycelia were washed with sterile distilled water and macerated thoroughly in liquid nitrogen, using a cooled pestle and mortar. The macerated mycelia (0.1 g) were immediately transferred into a micro-centrifuge tube (1.5 ml) and DNA was extracted using the DNA kit OmniPrep (G-Biosciences) and following the manufacturer’s instructions. The DNA was suspended in Tris-EDTA buffer (pH 8.0) and was treated with RNAse. The DNA quality and quantity were scored, using NanoDrop ND-1000.

#### Restriction fragment length polymorphism analysis using mitochondrial haplotyping and probe RG57

*Phytophthora* lineages are conventionally characterized by using mitochondrial haplotyping (Carter *et al.* 1990; Griffith & Shaw 1998) and restriction fragment length polymorphism (RFLP) analysis with the highly polymorphic probe RG57 (Goodwin *et al.* 1992). We used the same approach to compare *P. betacei* with previously reported lineages within the *Phytophthora* genus. The mitochondrial haplotype was determined using the PCR-RFLP method with reference strains US-1 and US-8 included as positive controls (Griffith & Shaw 1998; Ordoñez *et al.* 2000; Adler *et al.* 2002; Gavino & Fry 2002).

#### Strain typing using microsatellite markers

To assess the population differentiation among *P. infestans, P. betacei*, and *P. andina*, simple sequence repeats (SSRs) were analyzed, using the protocols developed previously by Lees *et al.* (2006) and described in the Eucablight Network’s Protocol section dated March 2008 (www.eucablight.org) (Table S2). A total of 116 *P. betacei* isolates obtained in this study (Table S1), 117 *P. infestans* and 17 *P. andina* isolates reported in Goss *et al.* (2014) were included in our analyses. Among the 117 *P. infestans* isolates, there were 17 distinct clonal lineages, as well as genotypically diverse isolates from Mexico and Northern Europe.

#### Population genetic analyses using microsatellite data

We used a principal component analysis (PCA) for the combined *P. betacei, P. infestans*, and *P. andina* microsatellite data. The allele frequencies at bi-allelic sites for the triploid *P. infestans* isolates (1/3 or 2/3) were unknown. To account for this uncertainty, we subsampled the alleles at each locus for each isolate. Since *adegenet* treats ploidy as a global parameter, we generated resampled datasets for the strains from all species, assuming that all the individuals across species had the same ploidy. To account for both the uncertainty in allele frequencies at bi-allelic sites in triploid *P. infestans* isolates and the fact that *adegenet* would require all samples to have the same ploidy, we generated 100 independent diploid and 100 independent triploid resampled datasets for the PCA (i.e. within each subsampled dataset, all individuals were diploids or triploids). One hundred diploid and one hundred triploid resampled datasets were created, and *adegenet* was run independently on each of them.

To estimate the number of populations that would best explain the genetic variance in the group of isolates studied, we used the Bayesian model-based clustering program STRUCTURE v2.3 (Pritchard *et al.* 2000). To account for allele frequency uncertainty at the bi-allelic triploid *P. infestans* loci and because ploidy is a global parameter in STRUCTURE, we used the same 200 resampled datasets described for the PCA. We ran STRUCTURE a total of 32,000 times: (2 ploidies) × (100 resampled datasets) × (8 populations, K = 1 to 8) × (20 repetitions for K selection). Each run involved 1,000,000 MCMC steps with a burn-in of 100,000 and used the following parameters: NOADMIX = 0, LINKAGE = 0, INFERALPHA = 1, ALPHA = 1.0, UNIFPRIORALPHA = 1, ALPHAMAX = 10.0, and FREQSCORR = 0. The *ΔK* method (Evanno *et al.* 2005) was used to infer the most likely number of clusters by evaluating the rate of change in the log probability of data between successive *K* values for each resampled dataset.

#### Strain typing using genotyping-by-sequencing

Genomic DNA was isolated with the DNeasy^®^ Plant Mini Kit (QIAGEN, Germany). Genotyping-by-sequencing (GBS) was performed (as described by Elshire *et al.* 2011) at the Institute of Genomic Diversity (Cornell University) for a total of 70 *Phytophthora* isolates. Among these, there were 12 *P. infestans* (10 from Colombia and two reference strains from the United States, US-8 and US-12), 35 *P. betacei* (clonal lineage EC-3), one *P. andina* (clonal lineage EC-3), five *P. andina* (clonal lineage EC-2), three *P. andina* isolates of unknown clonal lineage, five *P. mirabilis*, eight *P. ipomoeae*, and one *P. phaseoli* isolate (Table 1). Briefly, genome complexity was reduced by digesting the total genomic DNA from individual samples with the type II restriction endonuclease *Ape*KI, which recognizes a degenerate 5-bp sequence (GCWGC, where W is A or T) and creates a 5’ overhang (3 bp). The digested products were then ligated to adapter pairs with enzyme-compatible overhangs; one adapter contained the barcode sequence and the other the binding site for the Illumina sequencing primer. The GBS library fragment-size distributions were checked on a BioAnalyzer (Agilent Technologies, Inc., USA). The PCR products were quantified and diluted for sequencing on an Illumina HiSeq 2500 sequencer (Illumina Inc., USA). A 96-well plate, comprising 70 samples and one blank, was multiplexed on a single Illumina flow cell lane.

**Table 1.** Description of isolates of *P. infestans, P. andina*, and *P. betacei* used for the morphological, physiological, phylogenetic and host preference assays. Isolates used for SSR analysis are shown in Table S1

| Sample ID | Species | Locality<br>(Country/State/Locality) | Original<br>Host | Year | Locus<br>Lineage | Mitochondrial<br>Haplotype | Mating<br>Type | Assay<sup>a</sup> | Source |
| --- | --- | --- | --- | --- | --- | --- | --- | --- | --- |
| **1826** | *P. infestans* T-30-4 strain<sup>b</sup> | Scotland (SCRI) | NA | NA | NA | NA | NA | C | Grünwald lab culture collection |
| **4392** | *P. infestans* T-30-4 strain<sup>c</sup> | Scotland | NA | 2007 | NA | NA | NA | C | Grünwald lab culture collection |
| **US940494** | *P. infestans* | USA | *S. lycopersicum* | NA | US-12 | NA | A1 | C | Fry Lab culture collection |
| **US970001** | *P. infestans* | USA/Florida | *S. lycopersicum* | 1997 | US-17 | NA | A1 | A, B | Fry Lab culture collection |
| **US040009** | *P. infestans* | USA/New York | *S. tuberosum* | NA | US-8 | NA | A2 | A, B | Fry Lab culture collection |
| **US940480** | *P. infestans* | NA | NA | NA | US-8 | Ia | A2 | C | Fry Lab culture collection |
| **STG100** | *P. infestans* | Colombia/Nariño/Guachucal | *S. tuberosum* | 2013 | EC-1 | IIa | A1 | A, B | LAMFU<sup>d</sup> |
| **STT161** | *P. infestans* | Colombia/Nariño/Túquerres | *S. tuberosum* | 2013 | EC-1 | IIa | A1 | A, B | LAMFU<sup>d</sup> |
| **VPC7-10** | *P. infestans* | Colombia/Cundinamarca/Villa<br>Pinzon | *S. tuberosum* | 2015 | EC-1 | IIa | A1 | A, B | LAMFU<sup>d</sup> |
| **RC1-6** | *P. infestans* | Colombia/Cundinamarca/Rosal | *S. tuberosum* | 2015 | EC-1 | IIa | A1 | A, B | LAMFU<sup>d</sup> |
| **Z3-2** | *P. infestans* | Colombia/Cundinamarca/ Zipacón | *S. phureja* | 2007 | NA | NA | A1 | A, B, C, D | (Vargas *et al.* 2009) |
| **RB005** | *P. infestans* | Colombia/Nariño | *S. tuberosum* | 2013 | NA | NA | NA | C, D | LAMFU<sup>d</sup> |
| **RB003** | *P. infestans* | Colombia/Nariño | *S. tuberosum* | 2013 | NA | NA | NA | C | LAMFU<sup>d</sup> |
| **C003210** | *P. infestans* | Colombia/Nariño | *S. tuberosum* | 2013 | NA | NA | NA | C | LAMFU<sup>d</sup> |
| **C003B21** | *P. infestans* | Colombia/Nariño | *S. tuberosum* | 2013 | NA | NA | NA | C | LAMFU<sup>d</sup> |
| **PUA10096** | *P. infestans* | Colombia/Cundinamarca/Guasca | *S. phureja* | 2013 | NA | NA | A1 | C | LAMFU<sup>d</sup> |
| **C0008S** | *P. infestans* | Colombia/Putumayo/Sibundoy | *S. tuberosum* | 2013 | NA | NA | NA | C | LAMFU<sup>d</sup> |
| **SP03562** | *P. infestans* | Colombia/Putumayo/Sibundoy | *S. tuberosum* | 2013 | NA | NA | NA | C | LAMFU<sup>d</sup> |
| **EC 3399** | *P. andina*<sup>e</sup> | Ecuador | Anarrichomenum | NA | EC-2 | Ia | A2 | A, B, D | (Oliva *et al.* 2010; Goss *et al.* 2011) |
| **EC 3818** | *P. andina*<sup>e</sup> | Ecuador | Anarrichomenum | NA | EC-2 | Ia | A2 | A, B, D | (Oliva *et al.* 2010; Goss *et al.* 2011) |
| **EC 3510** | *P. andina*<sup>e</sup> | Ecuador | *S. betaceum* | NA | EC-3 | Ia | A1 | A, B, D | (Goss *et al.* 2011) |
| **EC 3836** | *P. andina*<sup>e</sup> | Ecuador | *S. betaceum* | 2008 | EC-3 | Ia | A1 | D | (Goss *et al.* 2011) |
| **EC 3780** | *P. andina*<sup>e</sup> | Ecuador | *S. hispidum* | NA | NA | Ic | NA | C | (Goss *et al.* 2011) |
| **EC 3818** | *P. andina*<sup>e</sup> | Ecuador | Anarrichomenum | NA | EC-2 | Ia | A2 | C | (Oliva *et al.* 2010; Goss *et al.* 2011) |
| **EC 3399** | *P. andina*<sup>e</sup> | Ecuador | Anarrichomenum | NA | EC-2 | Ia | A1 | C | (Oliva *et al.* 2010; Goss *et al.* 2011) |
| **EC 3821** | *P. andina*<sup>e</sup> | Ecuador | Anarrichomenum | NA | NA | Ia | NA | C | (Goss *et al.* 2011) |
| **EC 3189** | *P. andina*<sup>e</sup> | Ecuador | Anarrichomenum | NA | EC-2 | Ic | A2 | C | (Oliva *et al.* 2010; Goss *et al.* 2011) |
| **EC 3163** | *P. andina*<sup>e</sup> | Ecuador | Anarrichomenum | NA | EC-2 | Ic | A1 | C | (Oliva *et al.* 2010; Goss *et al.* 2011) |
| **EC 3678** | *P. andina*<sup>e</sup> | Ecuador | Anarrichomenum | NA | EC-2 | Ic | A1 | C | (Goss *et al.* 2011) |
| **EC 3563** | *P. andina*<sup>e</sup> | Ecuador | *S. quitoense* | NA | NA | Ia | A1 | C | (Goss *et al.* 2011) |
| **EC 3510** | *P. andina*<sup>e</sup> | Ecuador | *S. betaceum* | NA | EC-3 | Ia | A1 | C | Goss *et al.* 2011) |
| **MFN-N9022** | *P. betacei* | Colombia/ Nariño/Buesaco | *S. betaceum* | 2009 | EC-3 | Ia | A1 | A, B, C, D | This study |
| **MFN-N9039** | *P. betacei* | Colombia/Nariño/Buesaco | *S. betaceum* | 2009 | EC-3 | Ia | A1 | A, B, C | This study |
| **MFN-N9071** | *P. betacei* | Colombia/Nariño/ Iles | *S. betaceum* | 2009 | EC-3 | Ia | A1 | A, B, C | This study |
| **MFM-P8077** | *P. betacei* | Colombia/Putumayo/Colon | *S. betaceum* | 2008 | EC-3 | Ia | A1 | A, B, C | This study |
| **MFM-P8084** | *P. betacei* | Colombia/Putumayo/Colon | *S. betaceum* | 2008 | EC-3 | Ia | A1 | A, B, C, D | This study |
| **MFM-P8071** | *P. betacei* | Colombia/Putumayo/Colon | *S. betaceum* | 2008 | EC-3 | Ia | A1 | C | This study |
| **MFM-P8050** | *P. betacei* | Colombia/Putumayo/Colon | *S. betaceum* | 2008 | EC-3 | Ia | A1 | C | This study |
| **MFM-N9046** | *P. betacei* | Colombia/Nariño/ Pasto | *S. betaceum* | 2009 | EC-3 | Ia | A1 | C | This study |
| **MFM-N9057** | *P. betacei* | Colombia/Nariño/Consaca | *S. betaceum* | 2009 | EC-3 | Ia | A1 | C | This study |
| **MFM-N9056** | *P. betacei* | Colombia/Nariño/ Consaca | *S. betaceum* | 2009 | EC-3 | Ia | A1 | C | This study |
| **MFM-N9041** | *P. betacei* | Colombia/Nariño/ Consaca | *S. betaceum* | 2009 | EC-3 | Ia | A1 | C | This study |
| **MFM-P9127** | *P. betacei* | Colombia/Putumayo/ San Francisco | *S. betaceum* | 2009 | EC-3 | Ia | A1 | C | This study |
| **MFM-P8012** | *P. betacei* | Colombia/Putumayo/ Colon | *S. betaceum* | 2008 | EC-3 | Ia | A1 | C | This study |
| **MFM-P8064** | *P. betacei* | Colombia/Putumayo/ Santiago | *S. betaceum* | 2008 | EC-3 | Ia | A1 | C | This study |
| **MFM-P9146** | *P. betacei* | Colombia/Putumayo/ Sibundoy | *S. betaceum* | 2009 | EC-3 | Ia | A1 | C | This study |
| **MFM-P9128** | *P. betacei* | Colombia/Putumayo/ Sibundoy | *S. betaceum* | 2009 | EC-3 | Ia | A1 | C | This study |
| **MFM-N9025** | *P. betacei* | Colombia/Nariño/ Buesaco | *S. betaceum* | 2009 | EC-3 | Ia | A1 | C | This study; Forbes *et al.*, 2016 |
| **MFM-P9147** | *P. betacei* | Colombia/Putumayo/Sibundoy | *S. betaceum* | 2009 | EC-3 | Ia | A1 | C | This study |
| **MFM-P8075** | *P. betacei* | Colombia/Putumayo/ Colon | *S. betaceum* | 2008 | EC-3 | Ia | A1 | C | This study |
| **MFM-N9012** | *P. betacei* | Colombia/Nariño/ Buesaco | *S. betaceum* | 2009 | EC-3 | Ia | A1 | C | This study |
| **MFM-P8096** | *P. betacei* | Colombia/Putumayo/ San Francisco | *S. betaceum* | 2008 | EC-3 | Ia | A1 | C | This study |
| **MFM-P8099** | *P. betacei* | Colombia/Putumayo/ San Francisco | *S. betaceum* | 2008 | EC-3 | Ia | A1 | C | This study |
| **MFM-P9151** | *P. betacei* | Colombia/Putumayo/Sibundoy | *S. betaceum* | 2009 | EC-3 | Ia | A1 | C | This study |
| **MFM-P9153** | *P. betacei* | Colombia/Putumayo/Sibundoy | *S. betaceum* | 2009 | EC-3 | Ia | A1 | C | This study |
| **MFM-N9065** | *P. betacei* | Colombia/Nariño/ Iles | *S. betaceum* | 2009 | EC-3 | Ia | A1 | C | This study |
| **MFM-P9129** | *P. betacei* | Colombia/Putumayo/Sibundoy | *S. betaceum* | 2009 | EC-3 | Ia | A1 | C | This study |
| **S06298** | *P. betacei* | Colombia/Putumayo/ Santiago/ | *S. betaceum* | 2012 | EC-3 | Ia | A1 | C | This study |
| **MF-M-P8029** | *P. betacei* | Colombia/Nariño/ Buesaco | *S. betaceum* | 2008 | EC-3 | Ia | A1 | C | This study |
| **MF-M-P8093** | *P. betacei* | Colombia/Putumayo | *S. betaceum* | 2008 | EC-3 | Ia | A1 | C | This study |
| **MF-M-P9105** | *P. betacei* | Colombia/Putumayo/ San Francisco/ | *S. betaceum* | 2009 | EC-3 | Ia | A1 | C | This study |
| **S07198** | *P. betacei* | Colombia/Putumayo/ Sibundoy | *S. betaceum* | 2012 | EC-3 | Ia | A1 | C | This study |
| **CO1298** | *P. betacei* | Colombia/Putumayo/ Sibundoy | *S. betaceum* | 2012 | EC-3 | Ia | A1 | C | This study |
| **S00321** | *P. betacei* | Colombia/Putumayo/ Sibundoy | *S. betaceum* | 2012 | EC-3 | Ia | A1 | C | This study |
| **A01492** | *P. betacei* | Colombia/Antioquia | *S. betaceum* | 2012 | EC-3 | Ia | A1 | C | This study |
| **S07398** | *P. betacei* | Colombia/Putumayo/ Sibundoy | *S. betaceum* | 2012 | EC-3 | Ia | A1 | C | This study |
| **1260** | *P. mirabilis* | Mexico/Texcoco<sup>f</sup> | NA | NA | NA | NA | NA | C | Grünwald lab culture collection |
| **1991** | *P. mirabilis* | Mexico/ Coyoacán | *M. jalapa* | 2000 | NA | NA | NA | C | Grünwald lab culture collection |
| **1992** | *P. mirabilis* | Mexico/ Coyoacán | *M. jalapa* | 2000 | NA | NA | NA | C | Grünwald lab culture collection |
| **1276** | *P. mirabilis* | Mexico | *M. jalapa* | 1998 | NA | NA | NA | C | Grünwald lab culture collection |
| **1994** | *P. mirabilis* | Mexico/ Coyoacán | *M. jalapa* | 2007 | NA | NA | NA | C | Grünwald lab culture collection |
| **1271** | *P. ipomoeae* | Mexico | *I. longipedunculata* | 1999 | NA | NA | NA | C | Grünwald lab culture collection |
| **4667** | *P. ipomoeae* | Mexico | *I. longipedunculata* | 1999 | NA | NA | NA | C | Grünwald lab culture collection |
| **4666** | *P. ipomoeae* | Mexico | *I. longipedunculata* | 1999 | NA | NA | NA | C | Grünwald lab culture collection |
| **4669** | *P. ipomoeae* | Mexico | *I. longipedunculata* | 1999 | NA | NA | NA | C | Grünwald lab culture collection |
| **4670** | *P. ipomoeae* | Mexico | *I. longipedunculata* | 1999 | NA | NA | NA | C | Grünwald lab culture collection |
| **1270** | *P. ipomoeae* | Mexico | *I. longipedunculata* | 1999 | NA | NA | NA | C | Grünwald lab culture collection |
| **1989** | *P. ipomoeae* | Mexico | *I. longipedunculata* | 2000 | NA | NA | NA | C | Grünwald lab culture collection |
| **1990** | *P. ipomoeae* | Mexico/Michoacan | *Ipomoea* spp. | 1999 | NA | NA | NA | C | Grünwald lab culture collection |
| **5134** | *P. phaseoli* | USA | *Phaseolus lunatus* | 2003 | NA | NA | NA | C | Grünwald lab culture collection |

NA: data not available.
<sup>a</sup> Isolates used for A=morphological, B=Physiological, C=Phylogenetic analysis with GBS data, D=Host preference assay.
<sup>b</sup> F1 of a cross between two aggressive strains of *P. infestans* originally isolated from potato in the Netherlands. Isolate used for genome sequence.
<sup>c</sup> Duplicate of #1826
<sup>d</sup> LAMFU: The Mycology and Plant Pathology Laboratory at Universidad de los Andes, Bogotá, Colombia.
<sup>e</sup> Isolates classified as *P. andina* according to Oliva *et al.*, (2010).
<sup>f</sup> from CBS in the Netherlands, ATCC 64130

To sort each of the GBS barcode samples into separate fastq files, *Phytophthora* samples were demultiplexed using sabre (https://github.com/najoshi/sabre), allowing no mismatches within the barcode. In total, 1,992,701 tags were analyzed and mapped against the *P. infestans* T30-4 reference genome (Haas *et al.* 2009a), using Bowtie v2.2.3 (Langmead 2010). Out of this total number of reads, 917,890 (46.1%) were aligned to unique positions, 573,880 (28.8%) were aligned to multiple positions, and 500,931 (25.1%) could not be aligned to the reference genome. For SNP calling, each SAM sample alignment file was converted into a BAM file, followed by sorting and indexing, using SAMtools. SNPs and indels were called simultaneously, using the variant caller GATK v4.3.10 (McKenna *et al.* 2010a). The final dataset consisted of 70 samples with 23,480 SNPs obtained from GATK. To discard the presence of sequencing errors in our data, all samples that did not fulfill the following criteria were filtered out: mapping QUAL > 30, an overall coverage between 8 and 32X (cutoff values between 5% and 95% coverage), and a minor allele frequency (MAF) > 0.05.

#### Maximum likelihood phylogenetic analyses of SNP data

To infer the phylogenetic relationships of the *Phytophthora* 1c clade species (Table 1), we created a matrix where all high-quality SNP loci obtained from our GBS analyses were concatenated into a single alignment. We generated a maximum likelihood (ML) phylogenetic tree, using RAxML (Stamatakis 2006) under the general reversible nucleotide substitution model (GTR) with 1,000 bootstrap replicates to quantify branch support. The software jModelTest v. 2.1.7 was used to select the best-fit substitution model. *P. phaseoli* was used as an outgroup. The phylogenetic tree was drawn using Figtree v1.4.2 (http://tree.bio.ed.ac.uk/software/figtree/) (Rambaut 2009).

#### Population genetics analyses using genotyping-by-sequencing data

To corroborate the population structure analyses obtained by using 11 microsatellite loci, a PCA was conducted based on the 23,480 high-quality SNP markers obtained from our GBS analyses. High quality was defined as SNPs with MAF > 0.05 and with less than 20% missing data, that is, SNPs that were present in at least 80% of the strains assessed. Genetic structure was also estimated using the Bayesian assignment test implemented in the program STRUCTURE v2.3 (Pritchard *et al.* 2000) for high-quality SNP markers. These are defined as SNPs with MAF > 0.05 and with less than 10% missing data. We used a total of 48 samples: 12 for *P. infestans,* 29 for *P. betacei* and 7 for *P. andina* (EC-2). Run parameters were as follows: 24 runs with 4 repetitions with 100,000 MCMC steps and a burn-in period of 10,000 for 6 populations (K = 1 to 6), under the NOADMIX ancestry model and allele frequencies correlated. The ΔK of Evanno (Evanno *et al.* 2005) was calculated using the application Structure Harvester v. 0.6.94 (Earl & vonHoldt 2012) to infer the most likely number of clusters.

#### Whole genome sequencing and mitochondrial genome assembly

*Phytophthora betacei* strains were grown on liquid Plich medium for 10 to 15 days at 20°C for subsequent genomic DNA extraction as described above. Two *P. betacei* (P8084 and N9022) were sequenced using Illumina sequencing. A standard shotgun library (1x200bp) was constructed and sequenced by Beijing Genomic institute - BGI (HongKong, China) on an Illumina Hiseq2000 platform using paired-ends chemistry and 100 cycles. We generated 40 Gb of 96–100 bp paired-end reads from 2 libraries with insert lengths of 200 bp. We also generated 22 Gb of Illumina mate-pair libraries (6 kb insert size) for each of the *P. betacei* isolates. Read mapping was done with BWA-MEM 0.7.12 (Li 2013) with parameter k=10 using *P. infestans* as a reference (Haas *et al.* 2009). Variants were called with GATK 3.2-2 (McKenna *et al.* 2010b) using default parameters.

### Host pathogenicity assays

#### Evaluation of host preference

To estimate the effect of host specialization as a reproductive isolating barrier between *P. betacei* and *P. infestans*, we compared the fitness among these and *P. andina*, on different plant hosts. We used two isolates of each species for all infections assays: strains P8084 and N9022 for *P. betacei*, strains Z3-2 and RB005 for *P. infestans*, and strains EC3510 (EC-3, Ia) and EC3836 (EC-3, Ia) for *P. andina* all isolated from tree tomato (Table 1). We also included *P. andina* strains EC3399 (EC-2, Ia) and EC3818 (EC-2, Ia), isolated from hosts in the *Anarrichomenum* complex (Table 1). Each *Phytophthora* isolate was inoculated onto three different *Solanum* host species: *Solanum tuberosum* group *phureja* (yellow potato), *Solanum lycopersicum* (tomato), and *S. betaceum* variety Común (tree tomato). For isolates EC3399 and EC3818, no symptoms of infection were detected in inoculations done on *S. tuberosum, S. betaceum*, or *S. lycopersicum*. Thus, these isolates were excluded from the final analysis.

Plants were grown in the greenhouse (17 - 19°C), and leaves or leaflets were harvested after 8 to 10 weeks. Detached leaves were placed, abaxial side up, on the base of 90-mm petri plates containing moist paper towels. Three leaves were used per isolate as technical replicates. Each leaf was inoculated at four points with two 20-μl droplets of a sporangial suspension (3.5 × 10^4^ sporangia ml^−1^) on each side of the main vein. The petri plates were sealed with parafilm and incubated at 15 °C with a 16-h light period. Each experiment consisted of 4 hosts, 3 genotypes, 2 isolates per genotype, 3 leaflets per isolate, and 4 inoculation points per leaflet. The whole experiment was repeated three times.

The latent period, total lesion area, and number of sporangia produced were documented by taking daily pictures of the inoculated leaves from day 1 to day 9. The latent period was scored as the number of days it took from inoculations until sporangia were observed. The lesion area was scored as the necrotic area around the inoculation site, 9 days post inoculation (dpi), and was measured using Image J (rsb.info.nih.gov/ij/). The number of sporangia produced on each leaf 9 dpi was assessed by excising individual lesions and pooling them into 15-ml disposable polypropylene culture tubes with 3 ml of sterile distilled water. After vortexing for 10 sec sporangial numbers were counted at least twice using a haemocytometer. The total number of sporangia was calculated by averaging the sporangia counts per aliquot and then multiplying it by the dilution factor.

The total number of sporangia per day were calculated by dividing the total number of sporangia produced after 9 days by the number of days when sporangia were visible (9 days – latent period). The number of sporangia produced was calculated by subtracting the total number of sporangia produced 9 dpi from the sporangial concentration in the original inoculum suspension (2,800 sporangia per leaflet). We calculated fitness parameter for each replicate, as the reproductive rate of each genotype on each host as follows:

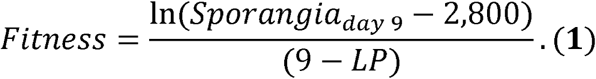

To quantify the variation in fitness, we fitted a full factorial, linear mixed model with the R package ‘nlme’ (function ‘lme’; Pinheiro *et al.* 2013). In the linear model, fitness was the response variable, genotype (*P. infestans, P. betacei*, or *P. andina*) and host (capiro potato, yellow potato, tomato, or tree tomato) were fixed effects, and strain (two independent isolates per genotype) was a random effect nested within a genotype. The significance of all interactions was assessed with Crawley’s (1993, 2002) ML approach, in which the full model containing all factors and interactions was fitted and then simplified by a series of stepwise deletions, starting with the fixed-effect interaction and progressing to the interaction terms. The critical probabilities for retaining factors and determining whether effects or interactions were significant were 5% for main effects and 1% for the two-way interactions. The linear model followed the formula:

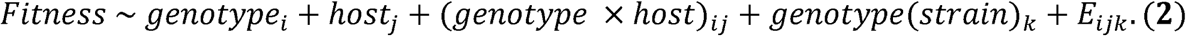

Because the residuals of this linear model were not normally distributed (Shapiro-Wilk normality test, W = 0.8342; P < 1 × 10^−15^) they were analyzed in a nonparametric framework. Fitness calculations on each host were compared among genotypes, using a Kruskal-Wallis test with the R package ‘stats’ (R Core team, 2013), followed by pairwise comparisons, using a Nemenyi test with a Tukey-Dist approximation for independent samples, using the R package ‘pmcmr’ (Pohlert 2014).

#### Phylogenetic analysis using mitochondrial genomes

We compared two newly assembled mitochondrial genomes of *P. betacei* sp. nov. (data taken from the whole genome sequences) and compared to data from Martin *et al.* (2016) in order to infer their phylogenetic position within clade 1c. MITObim (Hahn *et al.* 2013) was used to assembly the mitochondria of each genome. The process included 30 iterations using the quick approach with T30-4 haplotype Ia as the reference genome. Individual gene regions were aligned, using MAFFT v. 7.187 (Katoh & Toh 2010). Next, ML analyses were performed, using RAxML v.7.6.3 (Stamatakis 2006) as implemented on the CIPRES portal (Miller *et al.* 2010). The sequence alignment was partitioned into five subsets (rRNA genes, tRNA genes, first and second codon positions, third codon positions, and intergenic regions), similar to the work of Martin *et al.* (2016). The GTRGAMMA (GTR + G) model for nucleotide substitution was used but allowed the estimation of different shapes, GTR rates, and base frequencies for each partition. The majority rule criterion implemented in RAxML (-autoMRE) was used to assess clade support.

### Physiological and morphological characterization of *P. betacei*

#### Effect of temperature and culture media on colony morphology and mycelial radial growth

To assess the effect of temperature and culture media on colony morphology and on mycelial radial growth of *P. betacei* isolates (Table 1), we evaluated four distinct media and three different incubation temperatures. The four culture media tested were: V8 juice agar (V8), Potato Dextrose Agar (PDA, Oxoid Ltda, UK), Corn Meal Agar (CMA), and Tree Tomato Agar (TTA). For each medium, one agar plug (~ 5 mm diameter) of each actively growing culture was placed in the center of each petri plate (90 mm diameter). Colony morphology and mycelial radial growth for each isolate-media combination was evaluated 15 days post inoculation (dpi) by taking pictures using a Canon Digital EOS Rebel T3i / 600D camera (Tokyo, Japan). Colony morphology was described according to Erwin & Ribeiro (1996) and Gallegly & ChuanXue (2008). Radial growth was calculated by measuring the total mycelial growth area using ImageJ (rsb.info.nih.gov/ij/). To evaluate the optimum temperature for mycelial growth, all isolate-medium combinations previously described, were incubated at 4, 18, and 25°C in a dark chamber with constant humidity.

We followed the same procedure to evaluate colony morphology and mycelial radial growth in *P. andina* and *P. infestans.* Three isolates of *P. andina* EC3510 (EC-3; Ia), EC3399 (EC-2; Ia), and EC3818 (EC-2; Ia) and three isolates of *P. infestans* Z3-2 (EC-1), US040009 (US-8), and US970001 (US-17) (Table 1) were used. All combinations of isolates and media were tested in two independent blocks with two technical replicates per combination. Given the absence of mycelial growth at 4°C and 25°C, statistical analyses for mycelial radial growth were only conducted for isolates incubated at 18°C. For this analysis we compared data for *P. betacei, P. andina* (clonal lineage EC-2 and EC-3), and *P. infestans* for a total of 240 data points (1 temperature (18°C) × 15 isolates tested × 4 media × 2 technical replicates × 2 blocks). We assessed normality of the residuals of the linear models for each trait measured. In all cases, they were not normally distributed (Shapiro-Wilk test; P < 0.05; Table S2) and thus, we assessed whether there were differences in mycelial radial growth of the three species at different temperatures and different media.

We excluded the observations on PDA because *P. betacei* did not grow on this medium. We pooled the observations obtained from all other media and fitted a linear model where colony area was the response and the interaction between temperature and species was the only effect of the model. Pairwise comparisons were done using Tukey’s HSD (honest significant difference) test with the R library ‘multcomp’ (function ‘glht’).

#### Morphological characterization

We investigated whether isolates of *P. betacei* presented morphological differences with respect to isolates of *P. andina* and *P. infestans* (Table 1). We examined three morphological traits: *i*) sporangial morphology, *ii*) presence of hyphal swelling and chlamydospores, and *iii*) mycelia morphology as follows:

##### i Asexual reproductive structure (sporangia) morphology

We recorded the shape (length (μm), width (μm), and area (μm)), position, and caducity of sporangia on each isolate-medium combination described above, at 18°C (optimum growth temperature for all isolates; see results). Sporangia morphology was scored by measuring length (μm), width (μm), and area (μm) of sporangia for each isolate-medium combination. Mycelia from 15-day-old actively growing colony margins on each medium was excised and immersed directly in ~ 1 ml of sterile distilled water. From 10 to 30 sporangia were measured using a 60X oil objective and the FluoView FV1000 4.0 software (Olympus PlanApo 60X, 1.42NA) implemented in an Olympus IX81 microscope, for each isolate-medium combination. Pictures of sporangia were further analyzed and processed using the ImageJ software. Replicates were conducted for each isolate-medium combination in two separate blocks.

##### ii Presence of hyphal swelling and chlamydospores

Mycelia from 15-day-old actively growing colony margins on CMA medium (Difco), at 18°C, was excised and immersed directly in ~ 1 ml of sterile distilled water to score the presence of hyphal swelling and chlamydospores using a 60X oil objective in an Olympus IX81 microscope.

##### iii Mycelia morphology

The hyphal width (μm) of each of the three species on each of the four tested media (CMA, V8, PDA, and TTA) at 18°C were measured using mycelia from 15-day-old actively growing colony margins collected using a scalpel and immediately suspended in a drop (~ 50 μl) of sterile distilled water. Twenty randomly selected hyphae of each isolate-medium combination were measured using a 60X oil objective and the FluoView FV1000 4.0 software implemented in an Olympus IX81 microscope. Pictures were further analyzed and processed using the ImageJ software. Replicates were included for each combination in two separate blocks.

To assess heterogeneity among species in each of the studied traits, we used linear models where the measurements were the response variable and the species was the only fixed effect. We assessed the normality of the residuals of each linear model using a Shapiro-Wilk test (function ‘shapiro.test’, package ‘stats’; R Core team, 2013). Based on the Shapiro-Wilk test, residuals were not normally distributed in any of the linear models (Table S3, P< 0.05). Thus, we used the non-parametric Kruskal-Wallis test. To identify which group of isolates differed from each other, we performed multiple comparisons using non-parametric Nemenyi *post hoc* tests for Kruskal-Wallis. We performed all Kruskal-Wallis test using the R package ‘stats’ (R Core team, 2013) and the Nemenyi and Tukey post hoc tests using the Pairwise Multiple Comparison of Mean Ranks Package (‘pmcmr’) implemented in R (Pohlert 2014).

#### Discriminant analysis of morphological and physiological traits

Next, it was established whether the morphology of *P. betacei* and other *Phytophthora* species differed by visualizing all the morphological traits in a bidimensional plane using a discriminant function analysis (DA) based on the linear combination of morphological variables. To this end, a matrix with six traits (mycelial growth, hyphal width, sporangia length, sporangia width, sporangia area, and the sporangia length:width ratio) and a total of 15 individuals (five isolates for *P. betacei,* seven isolates for *P. infestans,* one isolate for *P. andina* clonal lineage EC-3, and two isolates for *P. andina* clonal lineage EC-2) was generated. Analyses were conducted using the “lda” function from the package ‘mass’ in R (Venables & Ripley 2002).

#### Molecular diagnosis of P. betacei based on SNP data

To distinguish *P. betacei, P. andina* (EC-2) and *P. infestans*, a set of 22,788 SNPs obtained from GBS data were analyzed for a total of 55 *Phytophthora* samples (12 *P. infestans*, 35 *P. betacei*, and 8 *P. andina* (EC-2); Table 1). Potentially diagnostic SNPs were selected calculating the allele frequencies and allele counts of each SNP for the entire dataset (55 samples and 22,788 SNPs). Major and minor alleles were obtained in each position. Samples belonging to each species were separated into three different files (*P. betacei, P. andina* (EC-2) and *P. infestans*) and allele counts were calculated for each dataset. SNPs with changes in the major allele in P. betacei were selected as candidates of differentiations relative to *P. infestans* and *P. andina* (EC-2) samples.

## Results

### Disease occurrence and *P. betacei* symptoms in the field

In 2008 and 2009, we identified a disease akin to late blight on tree tomato crops in southern Colombia. Field observations indicated that this disease can lead to the complete loss of the crop five to 10 days after the first symptoms are detected. In field, the pathogen is able to completely defoliate the tree in approximately one week. The symptoms of *P. betacei* on tree tomato differed from those generated by *P. infestans* on potato in forming concentric blighted areas that produced sporangia, and covered large areas of the leaves and petioles (Figure S2). In the field, no symptoms were observed on fruits, and the disease was rarely found on stems (Figure S2).

### Phylogenetic reconstruction and molecular population genetics

#### Mitochondrial haplotyping and RFLP analysis using probe RG57

All *P. betacei* isolates belonged to the Ia mitochondrial haplotype, and were assigned to the EC-3 clonal lineage based on the RG57 probe fingerprint pattern (Table 1).

#### Phylogenetic relationships of the Phytophthora 1c clade species using nuclear and mitochondrial genomes

A phylogenetic reconstruction using 23,480 nuclear SNPs showed *P. betacei, P. andina*, and *P. infestans* as more closely related to each another than to *P. ipomoeae* and *P. mirabilis* (Figure 1). Consistently, the former three species formed a monophyletic group. All *P. infestans* clonal lineages (EC-1, US-8, and US-12) formed a monophyletic group. *Phytophthora betacei* appeared as the sister group of the *P. andina* strains collected from wild *Solanaceae.* This *P. andina* group comprised isolates of the EC-2 clonal lineage with mitochondrial haplotypes Ia and Ic and some isolates of unknown clonal lineage. The two clades, *P. betacei* and *P. andina* (EC-2clonal lineage), were reciprocally monophyletic, providing evidence for the divergence of the two species. The only isolate of *P. andina* of the EC-3 clonal lineage that was included in our analysis grouped together with *P. betacei* (Figure 1). Interestingly, this strain was also isolated from *S. betaceum*.

**Figure 1.**
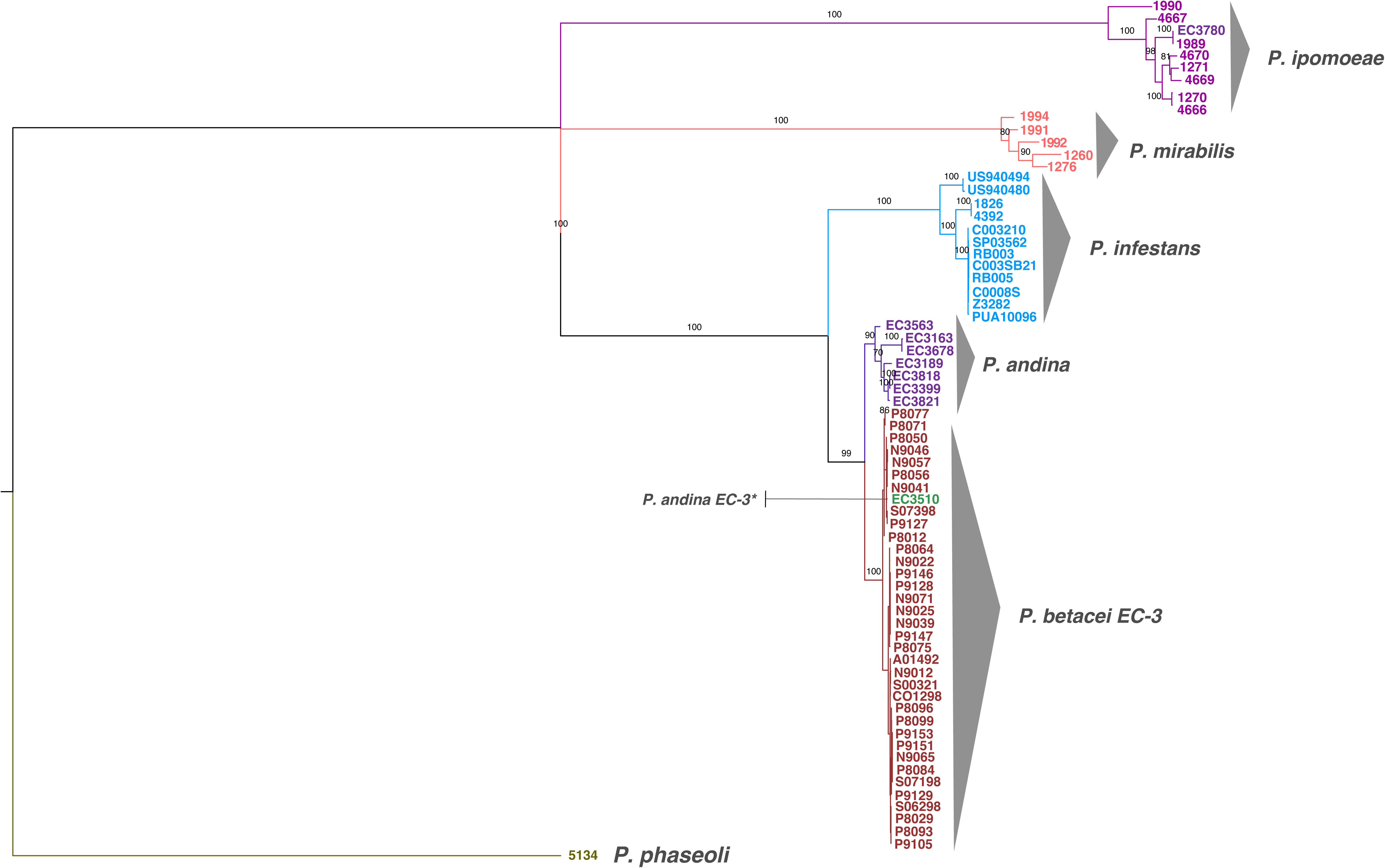
Phylogenetic relationship of *Phytophthora betacei* isolates and closely related species, based on genotyping by sequencing (GBS) data. The tree was inferred using Maximum Likelihood (ML) with *P. phaseoli* as an outgroup. Support values associated with branches correspond to ML bootstrap support values (BS). The three species, *P. infestans, P. betacei* and *P. andina* (clonal lineage EC-2) are clearly separated.

Phylogenetic analysis of the mitochondrial genome sequences did not differentiate among the three species of the *P. infestans sensu lato* complex, with one notable exception: *P. andina* clonal lineage EC-2 with the Ic mtDNA type appeared as the sister species of *P. mirabilis* (Figure 2).

**Figure 2.**
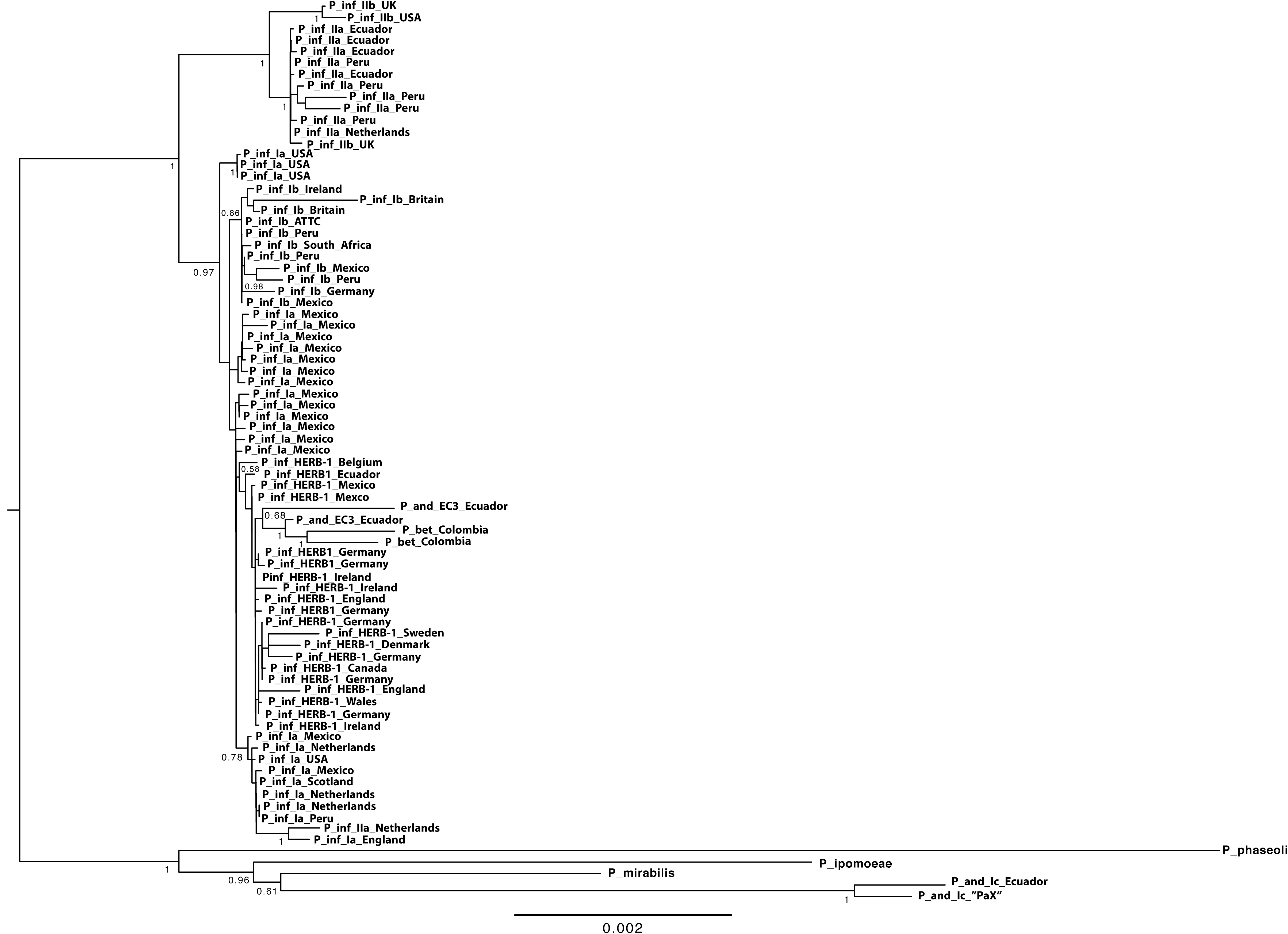
Maximum Likelihood (ML) phylogeny of complete mitochondrial genomes showing the hypothesized phylogenetic position of *P. betacei sp. nov*. Only bootstrap support values above 50 % are shown. **P_and** = *P. andina,* **P_inf** = *P. infestans*, **P_bet** = *P. betacei*. The analysis shows the polyphyletic nature of *P. andina* as currently described.

#### Population structure analyses

Microsatellite and whole-genome SNP data clearly differentiated among the populations of *P. infestans, P. betacei,* and *P. andina* (Figure 3). The results obtained from the PCA, using microsatellite data, suggested three genetic groups (Figure 3A). PC1 separated *P. infestans* and *P. betacei* (mean variance explained = 12.40%), indicating that a large proportion of the genetic variation is explained by the genetic differentiation between isolates belonging to these two species. PC2 (mean variance explained = 4.44%) showed intraspecific variation within *P. infestans*, which is larger than the variation within *P. betacei* or *P. andina*. Notably, the two *P. andina* strains of the EC-3 clonal lineage grouped closely with *P. betacei*. For this analysis, PCs 3, 4, and 5 primarily represent intraspecific variation within *P. infestans* (Figure S4). Similar results across all of the diploid and triploid resampled datasets suggested that regardless of the ploidy of the three species, the pattern of genetic differentiation is consistent (Figures S3 and S4, Table S6). A PCA on the GBS data including 23,480 SNPs supported the clustering of the SSR analysis (Figure 3B). The PCA shows strong genetic differentiation among strains of *P. infestans, P. betacei,* and *P. andina* (Figure 3B). PC1 accounted for 37.5% of the total variation and separated *P. infestans* and the *P. andina/P. betacei* clades. PC2 identified 5.8% of the variation between the strains and separated *P. betacei* and *P. andina* (Figure 3B).

**Figure 3.**
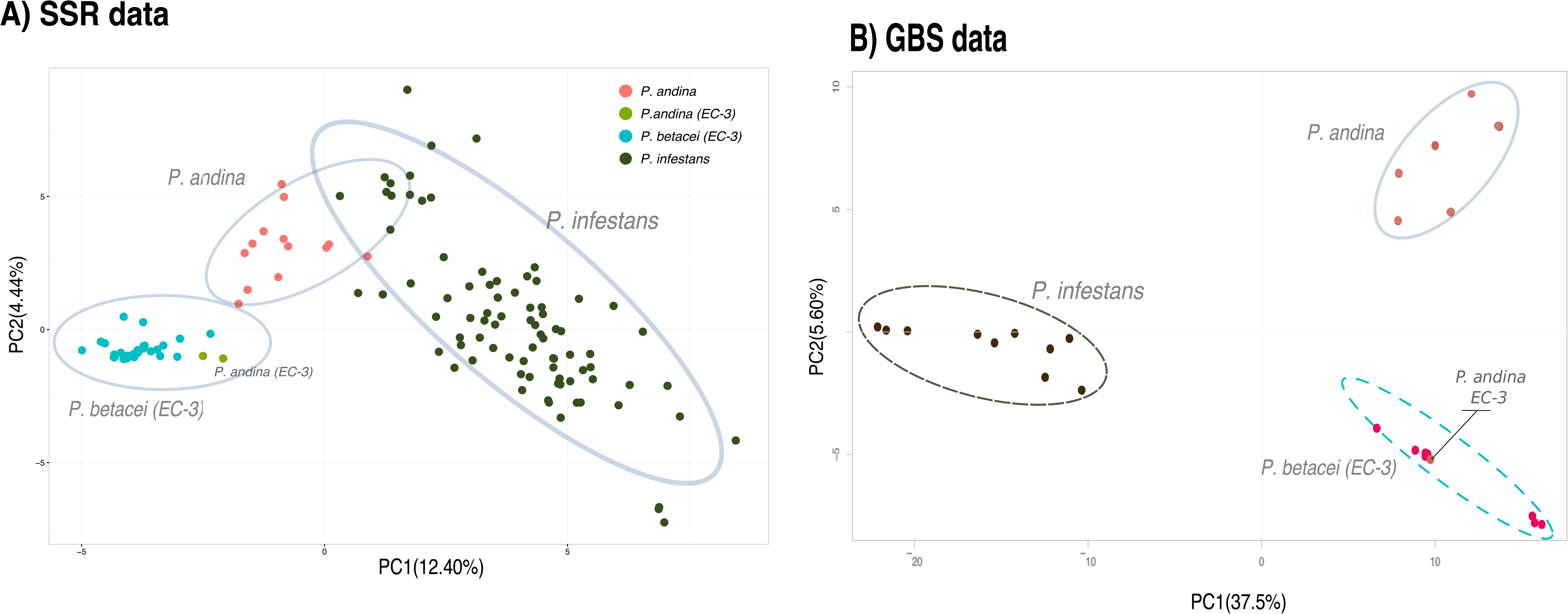
PCA analyses for microsatellite and GBS data showing the genetic structure for *P. betacei, P. andina* and *P. infestans*. **(A)** Results for microsatellite analyses. **(B)** Results for PCA analysis for GBS data.

We conducted two Bayesian assignment tests in STRUCTURE. For the GBS data, the most likely clustering was three populations (ΔK = 3) (Figure S5). The first genetic clusters matched with *P. infestans* sensu stricto and the second matched with *P. betacei*. The third cluster was assigned to *P. andina* samples (lineage EC-2) supporting the genetic differentiation between *P. infestans, P. andina* and *P. betacei* samples (Figure S5). For the SSR data, *Phytophthora andina* was most similar to *P. betacei* but contained genetic material from *P. infestans* (Figures S6-S8). Again, the two strains of *P. andina* of the EC-3 clonal lineage showed more genetic similarity to *P. betacei*. STRUCTURE for the SSR data revealed that the genetic variance in the sample was best explained by two genetic clusters’ populations (K = 2; Figure S6 and Tables S4 and S5). Population assignments for higher values of K are shown in Figures S6 and S7. The population assignments were robust to uncertainty in allele frequencies at bi-allelic triploid sites in *P. infestans* (K = 2 and equivalent individual assignments across diploid and triploid subsample datasets; Tables S4 and S5, and Figure S6 and S7). STRUCTURE analyses supports genetic differentiation between the three species.

### Host pathogenicity assays

The strongest line of evidence for a scenario of ecological speciation of plant pathogens comes from their host preference. Since *P. infestans, P. betacei*, and *P. andina* (EC-2) strains were isolated from different hosts, we tested the hypothesis that the three species were host specialized or had reduced fitness on their alternate host. We included isolates of *P. andina* of the EC-2 and EC-3 clonal lineages to make all possible pairwise comparisons. Isolates of *P. andina* of the EC-2 clonal lineage did not produce any symptoms on any of the hosts tested. *Phytophthora infestans* had higher fitness on tomato and yellow potato compared to *P. betacei* and *P. andina* (Table 2). *Phytophthora betacei* could not infect either tomato or potato but showed the highest fitness on tree tomato (Table 2). The *P. andina* (EC-3) strains assayed here were able to infect all hosts but showed lower fitness than *P. infestans* on three hosts (tomato, yellow potato, and tree tomato). They also displayed lower fitness on tree tomato compared to *P. betacei.* All pairwise comparisons indicated that strains of *P. infestans* and *P. betacei* displayed different fitness properties on every host assessed (Figure 4).

**Table 2.** **Pairwise comparisons of overall fitness on four different hosts.** Pairwise comparisons were made using a Kruskal Wallis rank sum test followed by a Nemenyi test with multiple comparisons. Statistically significant P-values (P < 0.05) are shown in bold.

| Host | Mean fitness (Standard Deviation) |  |  | Kruskal Wallis rank sum test |  | Nemenyi pairwise comparisons |  |  |
| --- | --- | --- | --- | --- | --- | --- | --- | --- |
| | *P. infestans* | *P. andina* (EC-3) | *P. betacei*<sup>a</sup> | $\chi^2$, df = 2 | P-value | *P. infestans* vs.<br>*P. andina* (EC-3) | *P. infestans* vs.<br>*P. betacei* | *P. betacei* vs. *P.*<br>*andina* (EC-3) |
| **Tomato**<br>(*S. lycopersicum*) | 0.0481<br>(±0.0589) | 0.0421 (±0.1055) | 0 (±0) | 18.0462 | $1.206 \times 10^{-4}$ | 0.5063 | **0.0097** | 0.2034 |
| **Potato** (*S. tuberosum*<br>*var. capiro*) | 0.0146 (±0.0341) | 0.0267 (±0.0486) | 0 (±0) | 4.7732 | 0.09194 | 0.89 | 0.76 | 0.45 |
| **Yellow potato**<br>(*S. phureja*) | 0.0604<br>(±0.0450) | 0.0545 (±0.0574) | 0 (±0) | 14.6133 | $6.711 \times 10^{-4}$ | 0.999 | **0.019** | **0.021** |
| **Tree tomato**<br>(*S. betaceum*) | 0.0659<br>(±0.0622) | 0.0298 (±0.0462) | 0.1025 (±0.0583) | 28.9611 | $5.143 \times 10^{-7}$ | **0.036** | **0.033** | **$5.7 \times 10^{-7}$** |

**Figure 4.**
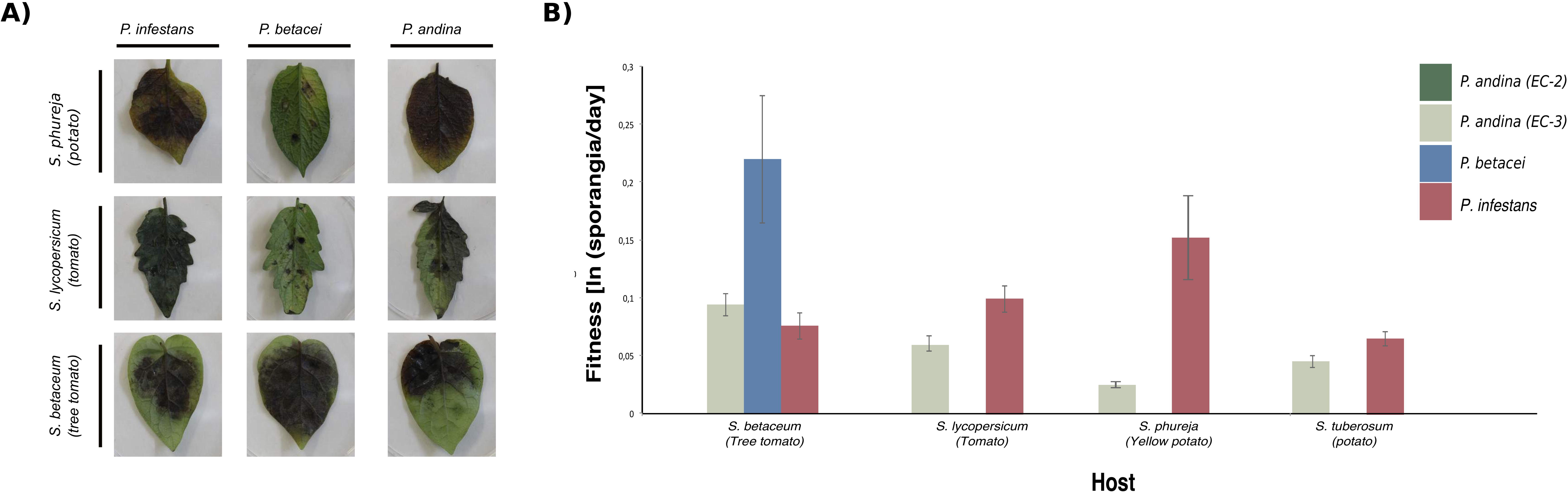
Host specialization induced strong premating reproductive isolation on *P. infestans, P. betacei* and *P. andina* isolates. **(A)** Fitness obtained from reciprocal infection assays. Results of infection for *P. infestans, P. betacei* and *P. andina* on the main host evaluated: potato (*S. tuberosum),* tomato (*S. lycopersicum)* and tree tomato (*S. betaceum)* after 9 days post inoculation (dpi). **(B)** Values of fitness (sporangia per day) for *P. infestans, P. betacei* and *P. andina*. On tree tomato, isolates of *P. betacei* showed significantly higher fitness than *P. infestans* and *P. andina*. Conversely, isolates of *P. betacei* are unable to infect other hosts where *P. infestans* thrives. Additionally, *P. andina* shows significantly lower fitness on all hosts tested.

### Discriminant analysis of morphological and physiological traits

*Phytophthora betacei* and *P. infestans* were highly differentiated when all morphological and physiological traits were analyzed jointly. Two variables (Table 3) explained 99% of the variance in a discriminant analysis (Figure 5). The first function (LD1) characterized the groups based mostly on the length:width ratio of sporangia (Table 3). For the second function (LD2), both hyphal width and length:width ratio helped discriminate among these groups of isolates (Table 3). Figure 5 shows the plot of the first and the second discriminant components for *P. betacei, P. infestans*, and *P. andina* (EC-2 and EC-3 clonal lineages). Physiological and morphological characterization of *P. betacei* is described in supplementary file 1.

**Table 3.** **Coefficients of the linear discriminant analysis (DA) for the quantitative morphological traits of *Phytophthora* isolates.** Values in bold represent the variables that best differentiate *P. betacei* from *P. infestans* and *P. andina*.

| Character | LD1 | LD2 |
| --- | --- | --- |
| Hyphal width (µm) | -1.0 | **-4.42** |
| Length of sporangia (µm) | 0.03 | -0.88 |
| Width of sporangia (µm) | -0.08 | 1.45 |
| Length:Width ratio | **-6.66** | **16.11** |
| Area of sporangia (µm) | 0.00 | 0.00 |
| Mycelial radial growth<br>(cm<sup>2</sup>) | -0.01 | 0.02 |

**Figure 5.**
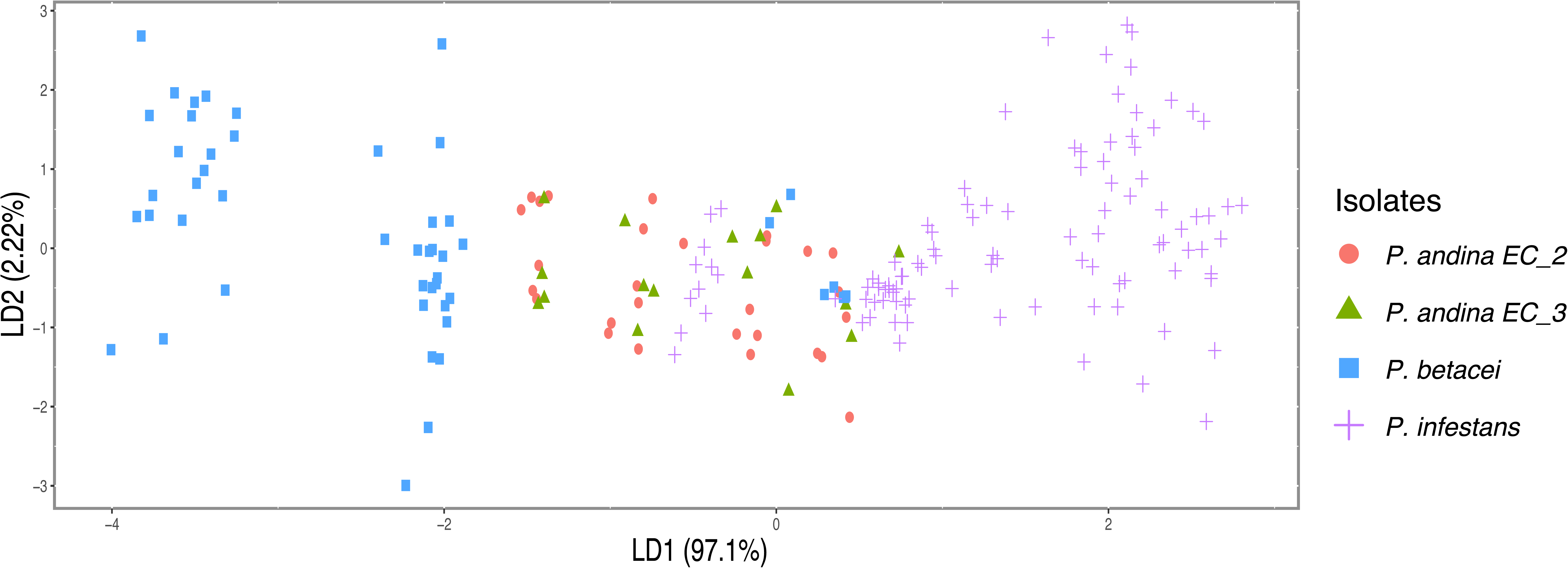
Plot of the morphological traits of *Phytophthora* species using a discriminant analysis (DA) showing the first and second discriminant components. *P. betacei* strains are shown in blue, *P. infestans* in green and *P. andina* in red.

### TAXONOMIC DESCRIPTION OF *Phytophthora betacei*

The taxonomic description of *P. betacei* was deposited in the MycoBank database (http://www.mycobank.org/) following standard taxonomy procedures.

***Phytophthora betacei* sp. nov**. M.F. Mideros, L.E. Lagos, et S. Restrepo, sp. nov. (Figure S9 - S11)

Mycobank Number No. **MB 815748**

#### Type material:— Holotype

Isolate of *P. betacei* from COLOMBIA, Putumayo, Colon, San Pedro locality, on infected leaves from *Solanum betaceum* (Solanaceae, Solanales), 1°13’26.9”N - 76°56’73.9”W, 24 Oct 2008, MF Mideros, (Andes-F 1172, holotype). ***Ex-type:*** LAMFU-COL-P8084

#### Description

*Phytophthora betacei* sp. nov. is an oomycete, plant pathogenic species that produced typical *Phytophthora* colonies that are white, smooth on V8 juice agar (V8). *P. betacei* grows well on V8 at 18°C. Aerial mycelia were generally abundant. Mycelial radial growth averaged 37.8 cm^2^ (SD = 9.6 cm^2^) after 15 days of incubation at 18°C. Sporangia of all isolates tested were borne terminally on the sporangiophore, and were caducous, ovoid, and semi-papillate with an average length of 36.3 μm (SD = 6.0 μm) and an average width of 17.3 μm (SD = 2.9 μm). The average length:width ratio was 2.1 (SD = 0.4). The area of sporangia was on average 436.3 μm^2^ (SD = 119.4 μm^2^). Growth of *P. betacei* isolates on potato dextrose agar (PDA) was limited. Mycelial radial growth was on average 0.27 cm^2^ (SD = 0.51 cm^2^) after 15 days of incubation at 18°C. No sporulation was detected on PDA. On corn meal agar (CMA), *P. betacei* isolates produced typical *Phytophthora* colonies that were white and smooth. All isolates were able to grow on CMA at 18°C. Aerial mycelium was generally abundant. Mycelial radial growth averaged 22.9 cm^2^ (SD ± 12.7 cm^2^) after 15 days of incubation at 18°C. Sporangia of all isolates were borne terminally on the sporangiophore, and were caducous, ovoid, and semi-papillate with an average length of 37.6 μm (SD = 5.4 μm) and an average width of 16.9 μm (SD = 2.3 μm). The average length:width ratio was 2.2 (SD = 0.2). The area of sporangia was on average 386.1 μm^2^ (SD = 33.1 μm^2^) on CMA. On tree tomato agar (TTA), *P. betacei* produced typical *Phytophthora* white smooth colonies. Mycelial radial growth and sporulation of *P. betacei* isolates was more abundant on TTA medium than on any of the other three media tested. Mycelial radial growth was on average 33.0 cm^2^ (SD = 6.2 cm^2^) after 15 days of incubation at 18°C. Sporangia of all isolates were borne terminally to the sporangiophore, and were caducous, ovoid, and semi-papillate with an average length of 39.3 μm (SD = 4.8 μm) and an average width of 15.8 μm (SD = 5.7 μm). The average length:width ratio was 2.6 (SD = 0.2). The area of sporangia was on average 311.5 μm^2^ (SD = 39.5 μm^2^). Hyphal swellings and chlamydospores were absent. Isolates were heterothallic with low oospore production (15.4 ± 10.9 oospores per mm^2^) and of abnormal appearance when crossed with a *P. infestans* strain of the A2 mating type (US040009). Oogonia were not ornamented. No self-fertile isolates were observed.

#### Material examined

Listed in Supplementary file 3.

#### Distribution

Isolates collected in southern Colombia, in the departments of Putumayo and Nariño.

#### Etymology

*“betacei”* refers to *S. betaceum*, the host plant from which the isolates were obtained.

#### Molecular diagnosis of P. betacei based on SNP data

From the GBS data, we identified a total of 22,788 SNPs from which 150 were able to discriminate *P. betacei* from *P. andina* (EC-2) and *P. infestans* samples. Although all 150 SNPs were classified as potentially diagnostic SNPs, this set of markers should then be validated in a larger *Phytophthora betacei* collection to validate their robustness to diagnose this species. All SNPs are listed in Supplementary file 3.

## Discussion

Here we describe the new species *P. betacei* which is closely related to *P. infestans* and *P. andina* but it is ecologically distinct. The divergence is recent but the levels of host specialization are very high suggesting ecological speciation in allopatry. The cross-pathogenicity tests showed strong host specificity when isolates of *P. betacei* were inoculated on *S. tuberosum* or *S. lycopersicum,* the main hosts of *P. infestans. Phytophthora infestans* is able to infect tree tomato, but its fitness (measured as the number of sporangia produced) on this host is relatively low compared with that on its more commonly described hosts (*S. tuberosum* and *S. lycopersicum*). Thus, host specificity might be playing an important role in maintaining gene flow between *P. infestans* and *P. betacei* restricted. Our findings provide novel insights into the evolutionary history of the Irish famine pathogen *P. infestans* and its close relatives. Furthermore, we also refine the species boundaries within the complex of *P. andina*, originally described as a polyphyletic taxon.

### *Phytophthora betacei* as a new species

We describe the new taxon, *P. betacei*, based on physiological, morphological, population genetic, and phylogenetic analyses, as well as differences in host specificity. All these analyses strongly support the designation of the new species *P. betacei* within the *Phytophthora* 1c clade.

The first line of evidence for the distinction between *P. betacei* and the other species of the *Phytophthora* 1c clade is the high genetic differentiation among the genetic groups. Nuclear phylogenies indicate that the triad *P. infestans, P. betacei,* and *P. andina* form a monophyletic clade whose closest known relatives are other members of the *Phytophthora* 1c clade (i.e., *P. ipomoeae, P. phaseoli*, and *P. mirabilis*). The three species, *P. betacei, P. infestans*, and *P. andina* (from the clonal lineage EC-2; see below) are clearly separated and *P. betacei* and *P. andina* (as defined here) are reciprocally monophyletic, suggesting that they lack recent gene flow and can be considered different species. Interestingly, the mitochondrial markers do not separate the three species. Our results are consistent with a scenario of speciation with secondary contact, and mitochondrial introgression, a phenomenon common across the tree of life (Funk & Omland 2003).

A second line of evidence for the existence of *P. betacei* as a separate species from *P. infestans sensu stricto* involves differences in allele frequencies in each of these genetic groups. All analyses using both SSR loci and SNP markers, suggest the existence of two discrete genetic clusters that correspond to *P. infestans sensu stricto* and *P. betacei. Phytophthora andina* has a less clear origin and this is discussed below. Our evidence suggests that *P. infestans* and *P. betacei* are isolated genetic groups with little detectable nuclear gene flow between them (Figure 3).

In addition to genetic variation, we determined whether *P. betacei* shows distinct morphological differences with *P. andina* and *P. infestans*. Four morphological (hyphal width and sporangial length, width, and length:width ratio) and one physiological (mycelial radial growth on four different media) traits were measured. The discriminant analysis clearly separated *P. betacei* from *P. infestans* in all media tested. The most striking morphological differences between *P. betacei* and *P. infestans* are the length:width ratio of sporangia, the hyphal width (μm), and the mycelial radial growth (cm^2^). Differences between *P. andina* and *P. infestans* or *P. betacei* are not as clear as the statistical differences between *P. andina* vs. *P. infestans* and *P. andina* vs. *P. betacei* and are dependent on the medium tested. Combining all the morphological variables, we show that strains of *P. betacei* collected in Colombia comprise a well-differentiated group of strains (Figure 5).

Our final and strongest line of evidence comes from infection assays on the native host range of the three species and from observations in nature. The host pathogenicity assays indicate that *P. betacei* is a tree tomato specialist unable to colonize potato and tomato (Figure 4). Conversely, *P. infestans* has low fitness on tree tomato, the only known host of *P. betacei*. These reciprocal differences in host pathogenicity represent a strong reproductive isolating mechanism between the two species. Host specificity is considered one of the most important isolating mechanisms between species of plant pathogens (reviewed in Harrington & Rizzo 1999; Coyne & Orr 2004). In asexual populations, host specialization could be associated with strong niche partition, which is common in species with asexual reproduction and strong local adaptation to the host (Poulin 2005; Halkett *et al.* 2006). Plant pathogens are commonly restricted to their hosts; thus, host specialization can result in a strong premating barrier (Stukenbrock 2013; Vialle *et al.* 2013).

Notably, we find that all isolates of *P. betacei* belonging to the EC-3 clonal lineage are closely related to the isolates previously described as *P. andina* EC-3 (Oliva *et al.* 2010; Goss *et al.* 2011, 2014; Lassiter *et al.* 2015; Martin *et al.* 2015). All EC-3 isolates form a monophyletic group differentiated from *P. infestans* and *P. andina* EC-2. We propose that all these EC-3 isolates should be considered *P. betacei* and all EC-2 isolates as *P.andina.* In this way, all species are rigorously defined as true monophyletic species.

### *Phytophthora andina* as a polyphyletic group

In the literature, *P. andina* has been reported to be polyphyletic and include the following three clonal lineages: *P. andina* EC-2 mitochondrial haplotype Ia, *P. andina* EC-2 mitochondrial haplotype Ic, and *P. andina* EC-3 (Adler *et al.* 2004; Gómez-Alpizar *et al.* 2007a). This species has been controversial since its erection since species are expected to be monophyletic with the expectation of descent from one common ancestor. *Phytophthora andina* was proposed to be a hybrid based on cloning nuclear haplotypes from several loci showing that one ancestor is *P. infestans* while the other ancestor remains to be described (Goss *et al.* 2011). Later, it has been hypothesized to have arisen from hybridization based on the conflicting phylogenetic information of mitochondrial and nuclear genealogies (Martin *et al.* 2015). Based on the hypothesis that *P. andina* was hybrid and the polyphyletic mitochondrial phylogenies in *P. andina,* we previously argued that *P. andina* was not appropriately described as a new species (Cárdenas *et al.* 2012). The identification of *P. betacei* as a new species, sheds some light on the origin of *P. andina*.

Our results support a monophyletic grouping of the EC-2 *P. andina* clonal lineages of mitochondrial haplotypes Ia and Ic that are closely related and form a monophyletic group distinct from *P. betacei* and *P. infestans.* Supported by phylogenetic and population genetic analyses, we suggest *P. andina* EC-3 should now be considered *P. betacei.* Our results indicate that the initial definition of *P. andina* included isolates that were either *P. betacei* or were closely related to *P. betacei,* namely, the EC-3 clonal lineage. At this point *P. betacei* cannot be considered a lineage of *P. andina* because it would reinforce the polyphyletic nature of *P. andina*. Again, a species cannot be described and proposed as polyphyletic. Our results showed the genetic and ecological separation of these two species, *P. betacei* and *P. andina* and our scenario of three species in the northern part of South America propose the most rigorous description of species.

Generally, our results confirm previous observations that *P. andina,* as currently described, is a polyphyletic group that requires redefinition (Gomez-Alpizar *et al.* 2008; Cárdenas *et al.* 2012; Forbes *et al.* 2012). Redefining *P. andina* including only strains of clonal lineage EC-2 makes this group monophyletic and provides a biologically rigorous species definition. We propose using *P. andina sensu lato* as the proper description of *P. andina* EC-2.

Whether there is reciprocal host specificity between *P. betacei* and *P. andina* (EC-2) was shown here. We have demonstrated that strains of *P. andina* of the EC-2 clonal lineage cannot infect *S. betaceum,* the only known host of *P. betacei.* Interestingly, isolates of the EC-3 clonal lineage have always been collected from *S. betaceum* plants, suggesting a strong isolating mechanism between *P. betacei* (clonal lineage EC-3) and *P. andina* (EC-2) in nature. Also, several authors have documented host specificity between *P. andina* EC-2 and EC-3 clonal lineages in nature (Adler *et al.* 2004; Gómez-Alpizar *et al.* 2007b; Oliva *et al.* 2010). All known EC-2 *P. andina* isolates have been collected in *Anarrichomenum* and other wild species, thus we hypothesize that the species might be specialized on these plants.

It is important to mention that a group of strains referred to as *P. andina* (new lineage PE-8) has recently been reported as infecting *S. betaceum* in Peru (Forbes *et al.* 2016). Further genetic, phylogenetic and population analyses and a greater number of isolates are needed to determine the identity, host range and fitness of isolates belonging to the PE-8 clonal lineage.

### Conclusions

We have provided several lines of evidence supporting the claim that *P. betacei* is a distinct, previously undescribed species within the *Phytophthora* 1c clade. Our findings and the scenario of the three species also resolve the polyphyletic nature of *P. andina*. The new species is ecologically distinct from its closely related species, *P. andina* and *P. infestans* showing high levels of host specialization suggesting ecological speciation in allopatry. The strong host specialization of *P. infestans* and *P. betacei* may act as premating barriers that restrict gene flow between these two species in nature. It remains unclear if host specialization facilitated or initiated the speciation process in the *P. infestans sensu lato* complex. However, in this report, we have demonstrated that ecological differences are important in the persistence of *P. infestans* and *P. betacei* as genetically isolated units across an overlapping area in the northern Andes. More studies are needed to further characterize the evolution of the closely related species and to understand the process of divergence in this group. In general, our results have implications for the understanding of how new plant pathogen species originate and persist. Our findings also highlight the importance of sampling plant pathogens of semi-domesticated or undomesticated hosts.

## Acknowledgements

This work was supported by the Department of Biological Sciences at Universidad de los Andes. Additional funding for this research was provided by the Research Fund of the School of Sciences and the Office of the Vice President for Research from Universidad de los Andes.

## SUPPLEMENTARY TABLE LEGENDS

**Table S1.** *Phytophthora betacei* isolates collected in this study and used for SSR analysis

| Collection number | Isolate number | Collection date | Location<br>(State/Municipality/Locality) | Geographical coordinates | Altitude (m) |
| --- | --- | --- | --- | --- | --- |
| ANDES-F 1081 | MFM-N9001 | 22/01/09 | Nariño/ Buesaco/ Vereda Medina Orejuela | 01°18.801"N 77°8.47,0"W | 2188 |
| ANDES-F 1082 | MFM-N9002 | 22/01/09 | Nariño/ Buesaco/ Vereda Medina Orejuela | 01°18.801"N 77°8.47,0"W | 2188 |
| ANDES-F 1083 | MFM-N9003 | 22/01/09 | Nariño/ Buesaco/ Vereda Medina Orejuela | 01°18.801"N 77°8.47,0"W | 2188 |
| ANDES-F 1084 | MFM-N9004 | 22/01/09 | Nariño/ Buesaco/ Vereda Medina Orejuela | 01°18.801"N 77°8.47,0"W | 2188 |
| ANDES-F 1085 | MFM-N9005 | 22/01/09 | Nariño/ Buesaco/ Vereda Medina Orejuela | 01°18.801"N 77°8.47,0"W | 2188 |
| ANDES-F 1086 | MFM-N9006 | 22/01/09 | Nariño/ Buesaco/ Vereda Medina Orejuela | 01°18.647"N 77°8.73,0"W | 2322 |
| ANDES-F 1087 | MFM-N9007 | 22/01/09 | Nariño/ Buesaco/ Vereda Medina Orejuela | 01°18.639"N 77°8.66,2"W | 2288 |
| ANDES-F 1088 | MFM-N9008 | 22/01/09 | Nariño/ Buesaco/ Vereda Medina Orejuela | 01°18.665"N 77°8.61,5"W | 2287 |
| ANDES-F 1089 | MFM-N9009 | 22/01/09 | Nariño/ Buesaco/ Vereda Medina Orejuela | 01°18.830"N 77°8.20,3"W | 2187 |
| ANDES-F 1090 | MFM-N9010 | 22/01/09 | Nariño/ Buesaco/ Vereda Medina Orejuela | 01°18.874"N 77°8.52,1"W | 2173 |
| ANDES-F 1091 | MFM-N9011 | 22/01/09 | Nariño/ Buesaco/ Vereda Medina Orejuela | 01°18.926"N 77°8.57,3"W | 2167 |
| ANDES-F 1092 | MFM-N9012 | 22/01/09 | Nariño/ Buesaco/ Vereda Medina Orejuela | 01°18.982"N 77°8.57,5"W | 2160 |
| ANDES-F 1093 | MFM-N9013 | 22/01/09 | Nariño/ Buesaco/ Vereda Medina Orejuela | 01°19.0,74"N 77°8.55,3"W | 2150 |
| ANDES-F 1094 | MFM-N9014 | 22/01/09 | Nariño/ Buesaco/ Vereda Medina Orejuela | 01°19.118"N 77°8.51,1"W | 2144 |
| ANDES-F 1095 | MFM-N9015 | 22/01/09 | Nariño/ Buesaco/ Vereda Medina Orejuela | 01°19.239"N 77°8.57,4"W | 2131 |
| **ANDES-F 1096** | MFM-N9016 | 22/01/09 | Nariño/ Buesaco/ Vereda Medina Orejuela | 01°19.231"N 77°8.56,0"W | 2136 |
| **ANDES-F 1097** | MFM-N9018 | 22/01/09 | Nariño/ Buesaco/ Vereda Medina Orejuela | 01°19.715"N 77°8.93,2"W | 2069 |
| **ANDES-F 1098** | MFM-N9019 | 22/01/09 | Nariño/ Buesaco/ Vereda Medina Orejuela | 01°19.930"N 77°9.0,36"W | 2107 |
| **ANDES-F 1099** | MFM-N9021 | 22/01/09 | Nariño/ Buesaco/ Vereda Llano Largo | 01°20.0,76"N 77°10.50,0"W | 2339 |
| **ANDES-F 1100** | MFM-N9022 | 22/01/09 | Nariño/ Buesaco/ Vereda Llano Largo | 01°20.23,9"N 77°10.49,8"W | 2303 |
| **ANDES-F 1101** | MFM-N9023 | 22/01/09 | Nariño/ Buesaco/ Vereda Llano Largo | 01°20.23,9"N 77°10.49,8"W | 2303 |
| **ANDES-F 1102** | MFM-N9024 | 22/01/09 | Nariño/ Buesaco/ Vereda Llano Largo | 01°20.23,9"N 77°10.49,10"W | 2303 |
| **ANDES-F 1103** | MFM-N9025 | 22/01/09 | Nariño/ Buesaco/ Vereda Llano Largo | 01°20.27,7"N 77°10.49,0"W | 2306 |
| **ANDES-F 1104** | MFM-N9027 | 22/01/09 | Nariño/ Buesaco/ Vereda Llano Largo | 01°20.27,7"N 77°10.49,0"W | 2306 |
| **ANDES-F 1105** | MFM-N9028 | 22/01/09 | Nariño/ Buesaco/ Vereda Llano Largo | 01°20.00,9"N 77°10.57,9 W | 2364 |
| **ANDES-F 1106** | MFM-N9029 | 22/01/09 | Nariño/ Buesaco/ Vereda Llano Largo | 01°20.02,6"N 77°10.53,8"W | 2343 |
| **ANDES-F 1107** | MFM-N9030 | 22/01/09 | Nariño/ Buesaco/ Vereda Llano Largo | 01°20.02,6"N 77°10.53,8"W | 2343 |
| **ANDES-F 1108** | MFM-N9031 | 22/01/09 | Nariño/ Buesaco/ Vereda Llano Largo | 01°19.59,5"N 77°11.53,8"W | 2180 |
| **ANDES-F 1109** | MFM-N9031 | 22/01/09 | Nariño/ Buesaco/ Vereda Llano Largo | 01°1.19"N 59,5°77.11"W | 2180 |
| **ANDES-F 1110** | MFM-N9033 | 02/02/09 | Nariño/ Buesaco/ Corregimiento Rosal del Monte | 01°17.29,0"N 77°10.42,6"W | NA |
| **ANDES-F 1111** | MFM-N9035 | 02/02/09 | Nariño/ Buesaco/ Corregimiento Rosal del Monte | 01°17.38,5"N 77°10.43,2"W | 2560 |
| **ANDES-F 1112** | MFM-N9036 | 02/02/09 | Nariño/ Buesaco/ Corregimiento Rosal del Monte | 01°17.38,7"N 77°10.42,3"W | 2543 |
| **ANDES-F 1113** | MFM-N9039 | 02/02/09 | Nariño/ Buesaco/ Corregimiento Rosal del Monte | 01°17.39,3"N 77°10.42,4"W | 2508 |
| **ANDES-F 1114** | MFM-N9041 | 02/02/09 | Nariño/ Buesaco/ Corregimiento Rosal del Monte | 01°17.39,3"N 77°10.42,4"W | 2520 |
| **ANDES-F 1115** | MFM-N9042 | 02/02/09 | Nariño/ Buesaco/ Corregimiento Rosal del Monte | 01°17.39,3"N 77°10.42,4"W | 2511 |
| **ANDES-F 1116** | MFM-N9046 | 20/02/09 | Nariño/ Pasto/ Corregimiento de Mocondino | 01°11.9"N 77°14.8,9"W | 2796 |
| **ANDES-F 1117** | MFM-N9056 | 25/03/09 | Nariño/ Consaca/ Vereda el Tejar | 01°12.52,74"N 77°27.33,7"W | 1861 |
| **ANDES-F 1118** | MFM-N9057 | 25/03/09 | Nariño/ Consaca/ Vereda el Tejar | 01°12.52,8"N 77°27.33,6"W | 1892 |
| **ANDES-F 1119** | MFM-N9065 | 26/03/09 | Nariño/ Iles/ Vereda Villa Nueva | 00°58.41,1"N 77°32.4"W | 2938 |
| **ANDES-F 1120** | MFM-N9071 | 26/03/09 | Nariño/ Iles/ Vereda Villa Nueva | 00°58.41,1"N 77°32.4"W | 2938 |
| **ANDES-F 1127** | MFM-P8010 | 14/09/08 | Putumayo/ Colon/ Casco Urbano | 01°11.0,94"N 76°58.35,4"W | 2107 |
| **ANDES-F 1128** | MFM-P8011 | 14/09/08 | Putumayo/ Colon/ Casco Urbano | 01°11.0,98"N 76°58.22,5"W | 2093 |
| **ANDES-F 1129** | MFM-P8012 | 14/09/08 | Putumayo/ Colon/ Casco Urbano | 01°11.6"N 76°58.22,8"W | 2094 |
| **ANDES-F 1143** | MFM-P8049 | 04/10/08 | Putumayo/ Colon/ Casco Urbano | 01°11.750"N 76°58.33,7"W | 2098 |
| **ANDES-F 1144** | MFM-P8050 | 04/10/08 | Putumayo/ Colon/ Casco Urbano | 01°11.750"N 76°58.33,7"W | 2098 |
| **ANDES-F 1145** | MFM-P8051 | 04/10/08 | Putumayo/ Colon/ Casco Urbano | 01°11.768"N 76°58.36,3"W | 2130 |
| **ANDES-F 1170** | MFM-P8082 | 13/10/08 | Putumayo/ Colon/ Vereda La Josefina | 01°10.802"N 76°58.89,9"W | 2090 |
| **ANDES-F 1157** | MFM-P8069 | 11/10/08 | Putumayo/ Colon/ Vereda Las Palmas | 01°11.750"N 76°57.45,0"W | NA |
| **ANDES-F 1158** | MFM-P8070 | 11/10/08 | Putumayo/ Colon/ Vereda Las Palmas | 01°11.765"N 76°57.45,6"W | NA |
| **ANDES-F 1159** | MFM-P8071 | 11/10/08 | Putumayo/ Colon/ Vereda Las Palmas | 01°11.967"N 76°57.54,3"W | 2107 |
| **ANDES-F 1160** | MFM-P8072 | 11/10/08 | Putumayo/ Colon/ Vereda Las Palmas | 01°12.0,009"N 76°57.57,8"W | NA |
| **ANDES-F 1161** | MFM-P8073 | 11/10/08 | Putumayo/ Colon/ Vereda Las Palmas | 01°12.0,009"N 76°57.57,8"W | NA |
| **ANDES-F 1162** | MFM-P8074 | 11/10/08 | Putumayo/ Colon/ Vereda San Pedro | 01°13.200"N 76°56.57,3"W | 2220 |
| **ANDES-F 1163** | MFM-P8075 | 11/10/08 | Putumayo/ Colon/ Vereda San Pedro | 01°13.100"N 76°56.45,5"W | 2168 |
| **ANDES-F 1164** | MFM-P8076 | 11/10/08 | Putumayo/ Colon/ Vereda San Pedro | 01°13.0,5"N 76°56.47,5"W | 2167 |
| **ANDES-F 1165** | MFM-P8077 | 11/10/08 | Putumayo/ Colon/ Vereda San Pedro | 01°12.949"N 76°56.49,4"W | 2151 |
| **ANDES-F 1166** | MFM-P8078 | 11/10/08 | Putumayo/ Colon/ Vereda San Pedro | 01°12.957"N 76°56.53,8"W | 2156 |
| **ANDES-F 1167** | MFM-P8079 | 11/10/08 | Putumayo/ Colon/ Vereda San Pedro | 01°12.912"N 76°56.50,0"W | 2153 |
| **ANDES-F 1168** | MFM-P8080 | 11/10/08 | Putumayo/ Colon/ Vereda San Pedro | 01°12.885"N 76°56.57,2"W | 2148 |
| **ANDES-F 1171** | MFM-P8083 | 24/10/08 | Putumayo/ Colon/ Vereda San Pedro | 01°13.269"N 76°56.73,9"W | 2305 |
| **ANDES-F 1172** | MFM-P8084 | 24/10/08 | Putumayo/ Colon/ Vereda San Pedro | 01°13.269"N 76°56.73,9"W | 2305 |
| **ANDES-F 1175** | MFM-P8087 | 07/11/08 | Putumayo/ San Francisco/ Casco Urbano | 01°10.629"N 76°52.69,4"W | 2148 |
| **ANDES-F 1176** | MFM-P8088 | 07/11/08 | Putumayo/ San Francisco/ Casco Urbano | 01°10.629"N 76°52.69,4"W | 2148 |
| **ANDES-F 1177** | MFM-P8091 | 07/11/08 | Putumayo/ San Francisco/ Casco Urbano | 01°10.550"N 76°52.49,4"W | 2220 |
| **ANDES-F 1193** | MFM-P9115 | 17/01/09 | Putumayo/ San Francisco/ Casco Urbano | 01°10.629"N 76°52.69,4"W | 2148 |
| **ANDES-F 1178** | MFM-P8093 | 07/11/08 | Putumayo/ San Francisco/ Vereda Chinayaco | 01°9.426"N 76°54.22,3"W | 2150 |
| **ANDES-F 1179** | MFM-P8094 | 07/11/08 | Putumayo/ San Francisco/ Vereda Chinayaco | 01°9.426"N 76°54.22,3"W | 2150 |
| **ANDES-F 1180** | MFM-P8095 | 07/11/08 | Putumayo/ San Francisco/ Vereda Chinayaco | 01°9.319"N 76°54.21,2"W | 2064 |
| **ANDES-F 1181** | MFM-P8096 | 07/11/08 | Putumayo/ San Francisco/ Vereda Chinayaco | 01°9.319"N 76°54.21,2"W | 2064 |
| **ANDES-F 1182** | MFM-P8097 | 07/11/08 | Putumayo/ San Francisco/ Vereda Chinayaco | 01°9.435"N 76°54.15,3"W | 2135 |
| **ANDES-F 1183** | MFM-P8098 | 07/11/08 | Putumayo/ San Francisco/ Vereda Chinayaco | 01°9.450"N 76°54.10,0"W | 2142 |
| **ANDES-F 1184** | MFM-P8099 | 07/11/08 | Putumayo/ San Francisco/ Vereda Chinayaco | 01°9.531"N 76°54.0,63"W | 2115 |
| **ANDES-F 1186** | MFM-P9104 | 17/01/09 | Putumayo/ San Francisco/ Vereda La Menta | 01°9.407"N 76°56.42,6"W | 2084 |
| **ANDES-F 1187** | MFM-P9105 | 17/01/09 | Putumayo/ San Francisco/ Vereda La Menta | 01°9.327"N 76°56.52,2"W | 2085 |
| **ANDES-F 1188** | MFM-P9108 | 17/01/09 | Putumayo/ San Francisco/ Vereda La Menta | 01°9.8,7"N 76°57.33,3"W | 2087 |
| **ANDES-F 1189** | MFM-P9109 | 17/01/09 | Putumayo/ San Francisco/ Vereda La Menta | 01°9.9,8"N 76°57.16,0"W | 2085 |
| **ANDES-F 1190** | MFM-P9110 | 17/01/09 | Putumayo/ San Francisco/ Vereda La Menta | 01°9.9,4"N 76°57.17,3"W | 2086 |
| **ANDES-F 1191** | MFM-P9113 | 17/01/09 | Putumayo/ San Francisco/ Vereda San Antonio de Porotoyaco | 01°9.6,8"N 76°54.27,3"W | 2152 |
| **ANDES-F 1192** | MFM-P9114 | 17/01/09 | Putumayo/ San Francisco/ Vereda San Antonio de Porotoyaco | 01°9.6,8"N 76°54.27,3"W | 2151 |
| **ANDES-F 1195** | MFM-P9127 | 22/01/09 | Putumayo/ San Francisco/ Vereda San Jose del Chunga | 01°7.51,3"N 76°58.0,8"W | 2457 |
| **ANDES-F 1185** | MFM-P9102 | 17/01/09 | Putumayo/ San Francisco/ Vereda San Silvestre | 01°10.2,2"N 76°56.21,6"W | 2101 |
| **ANDES-F 1194** | MFM-P9120 | 22/01/09 | Putumayo/ San Francisco/ Vereda San Silvestre | 01°10.4,2"N 76°56.22,5"W | 2086 |
| **ANDES-F 1130** | MFM-P8014 | 13/09/08 | Putumayo/ Santiago/ without precise locality | 01°6.56,8"N 76°59.10,7"W | 2116 |
| **ANDES-F 1131** | MFM-P8015 | 13/09/08 | Putumayo/ Santiago/ without precise locality | 01°6.57,6"N 76°59.0,92"W | 2115 |
| **ANDES-F 1132** | MFM-P8016 | 12/09/08 | Putumayo/ Santiago/ without precise locality | 01°7.3,7"N 76°59.11,5"W | 2096 |
| **ANDES-F 1133** | MFM-P8017 | 13/09/08 | Putumayo/ Santiago/ without precise locality | 01°7.0,63"N 76°59.14,4"W | 2093 |
| **ANDES-F 1134** | MFM-P8018 | 13/09/08 | Putumayo/ Santiago/ without precise locality | 01°7.3,7"N 76°59.11,5"W | 2096 |
| **ANDES-F 1125** | MFM-P8006 | 13/09/08 | Putumayo/ Santiago/ Vereda Balsayaco | 01°6.788"N 76°58.65,4"W | 2095 |
| **ANDES-F 1126** | MFM-P8008 | 13/09/08 | Putumayo/ Santiago/ Vereda Balsayaco | 01°6.949"N 76°58.61,9"W | 2078 |
| **ANDES-F 1136** | MFM-P8028 | 26/09/08 | Putumayo/ Santiago/ Vereda Balsayaco | 01°6.58,9"N 76°58.44,9"W | 2093 |
| **ANDES-F 1141** | MFM-P8034 | 26/09/08 | Putumayo/ Santiago/ Vereda Balsayaco | 01°6.58,9"N 76°58.44,9"W | 2093 |
| **ANDES-F 1123** | MFM-P8004 | 08/09/08 | Putumayo/ Santiago/ Vereda Cascajo | 01°7.820"N 77°1.30,0"W | 2246 |
| **ANDES-F 1124** | MFM-P8005 | 08/09/08 | Putumayo/ Santiago/ Vereda Cascajo | 01°7.846"N 77°1.16,0"W | 2247 |
| **ANDES-F 1169** | MFM-P8081 | 13/10/08 | Putumayo/ Santiago/ Vereda El Diviso | 01°9.312"N 76°59.72,5"W | 2087 |
| **ANDES-F 1121** | MFM-P8001 | 08/09/08 | Putumayo/ Santiago/ Vereda Muchivioy | 01°7.381"N 77°0.61,6"W | 2296 |
| **ANDES-F 1122** | MFM-P8003 | 08/09/08 | Putumayo/ Santiago/ Vereda Muchivioy | 01°7.542"N 77°7.50,1"W | 2310 |
| **ANDES-F 1137** | MFM-P8029 | 26/09/08 | Putumayo/ Santiago/ Vereda Quinchoa Pamba | 01°8.37,8"N 76°59.35,5"W | 2090 |
| **ANDES-F 1138** | MFM-P8030 | 27/09/08 | Putumayo/ Santiago/ Vereda Quinchoa Pamba | 01°8.4,4"N 76°59.20,8"W | 2088 |
| **ANDES-F 1139** | MFM-P8031 | 27/09/08 | Putumayo/ Santiago/ Vereda Quinchoa Pamba | 01°8.4,4"N 76°59.20,8"W | 2088 |
| **ANDES-F 1140** | MFM-P8032 | 27/09/08 | Putumayo/ Santiago/ Vereda Quinchoa Pamba | 01°8.57,5"N 76°59.45,2"W | 2088 |
| **ANDES-F 1135** | MFM-P8022 | 26/09/08 | Putumayo/ Santiago/ Vereda San Andres | 01°7.35,2"N 76°59.23,6"W | 2102 |
| **ANDES-F 1142** | MFM-P8047 | 04/10/08 | Putumayo/ Santiago/ Vereda Vichoy | 01°10.563"N 76°58.91,2"W | 2085 |
| **ANDES-F 1146** | MFM-P8054 | 04/10/08 | Putumayo/ Santiago/ Vereda Vichoy | 01°10.651"N 76°59.38,9"W | 2152 |
| **ANDES-F 1147** | MFM-P8055 | 04/10/08 | Putumayo/ Santiago/ Vereda Vichoy | 01°10.620" N 76°59.42,7"W | 2152 |
| **ANDES-F 1148** | MFM-P8056 | 04/10/08 | Putumayo/ Santiago/ Vereda Vichoy | 01°10.542"N 76°59.40,6"W | 2132 |
| **ANDES-F 1149** | MFM-P8057 | 04/10/08 | Putumayo/ Santiago/ Vereda Vichoy | 01°10.733"N 76°59.72,2"W | 2175 |
| **ANDES-F 1150** | MFM-P8058 | 04/10/08 | Putumayo/ Santiago/ Vereda Vichoy | 01°10.540"N 76°59.69,8"W | 2165 |
| **ANDES-F 1151** | MFM-P8059 | 04/10/08 | Putumayo/ Santiago/ Vereda Vichoy | 01°10.533"N 76°59.70,1"W | 2157 |
| **ANDES-F 1152** | MFM-P8060 | 04/10/08 | Putumayo/ Santiago/ Vereda Vichoy | 01°10.404"N 76°59.58,8"W | NA |
| **ANDES-F 1153** | MFM-P8061 | 04/10/08 | Putumayo/ Santiago/ Vereda Vichoy | 01°10.404"N 76°59.58,8"W | NA |
| **ANDES-F 1154** | MFM-P8063 | 05/10/08 | Putumayo/ Santiago/ Vereda Vichoy | 01°10.294"N 76°59.36,8"W | 2102 |
| **ANDES-F 1155** | MFM-P8064 | 05/10/08 | Putumayo/ Santiago/ Vereda Vichoy | 01°10.257"N 76°59.41,3"W | 2100 |
| **ANDES-F 1156** | MFM-P8066 | 05/10/08 | Putumayo/ Santiago/ Vereda Vichoy | 01°10.157"N 76°59.52,8"W | 2101 |
| **ANDES-F 1173** | MFM-P8085 | 24/10/08 | Putumayo/ Santiago/ Vereda Vichoy | 01°10.769"N 76°58.88,8"W | 2085 |
| **ANDES-F 1174** | MFM-P8086 | 24/10/08 | Putumayo/ Santiago/ Vereda Vichoy | 01°9.876"N 76°59.69,2"W | 2116 |
| **ANDES-F 1202** | MFM-P9150 | 05/02/09 | Putumayo/ Sibundoy/ Casco Urbano | 01°11.58,3"N 76°54.38,1"W | 2142 |
| **ANDES-F 1204** | MFM-P9153 | 05/02/09 | Putumayo/ Sibundoy/ Casco Urbano | 01°12.31,5"N 76°55.24,9"W | 2151 |
| **ANDES-F 1199** | MFM-P9146 | 05/02/09 | Putumayo/ Sibundoy/ Vereda Cabuyayaco | 01°11.17,3"N 76°56.11,0"W | 2096 |
| **ANDES-F 1196** | MFM-P9128 | 22/01/09 | Putumayo/ Sibundoy/ Vereda El Ejido | 01°11.22,5"N 76°56.7,4"W | 2081 |
| **ANDES-F 1197** | MFM-P9129 | 22/01/09 | Putumayo/ Sibundoy/ Vereda Machindinoy | 01°7.54,8"N 76°55.25,9"W | 2108 |
| **ANDES-F 1198** | MFM-P9144 | 05/02/09 | Putumayo/ Sibundoy/ Vereda San Felix | 01°10.50,7"N 76°56.16,8"W | 2087 |
| **ANDES-F 1200** | MFM-P9147 | 05/02/09 | Putumayo/ Sibundoy/ Vereda San Felix | 01°10.49,1"N 76°55.41,3"W | 2107 |
| **ANDES-F 1201** | MFM-P9148 | 05/02/09 | Putumayo/ Sibundoy/ Vereda San Felix | 01°11.19,4"N 76°55.36,0"W | 2088 |
| **ANDES-F 1203** | MFM-P9151 | 05/02/09 | Putumayo/ Sibundoy/ Vereda San Felix | 01°11.22,8"N 76°55.41,3"W | 2092 |
| **ANDES-F 1205** | MFM-P9154 | 05/02/09 | Putumayo/ Sibundoy/ Vereda San Felix | 01°11.04,0"N 76°56.10,6"W | 2097 |
| **ANDES-F 1206** | MFM-P9158 | 05/02/09 | Putumayo/ Sibundoy/ Vereda San Felix | 01°10.34,1"N 76°55.0,35"W | 2104 |
| **ANDES-F 1208** | MFM-P9164 | 06/02/09 | Putumayo/ Sibundoy/ Vereda San Felix | 01°12.41,3"N 76°55.10,6"W | 2201 |
| **ANDES-F 1207** | MFM-P9159 | 06/02/09 | Putumayo/ Sibundoy/ Vereda Villaflores | 01°12.38,3"N 76°55.5"W | 2199 |

**Table S2.**
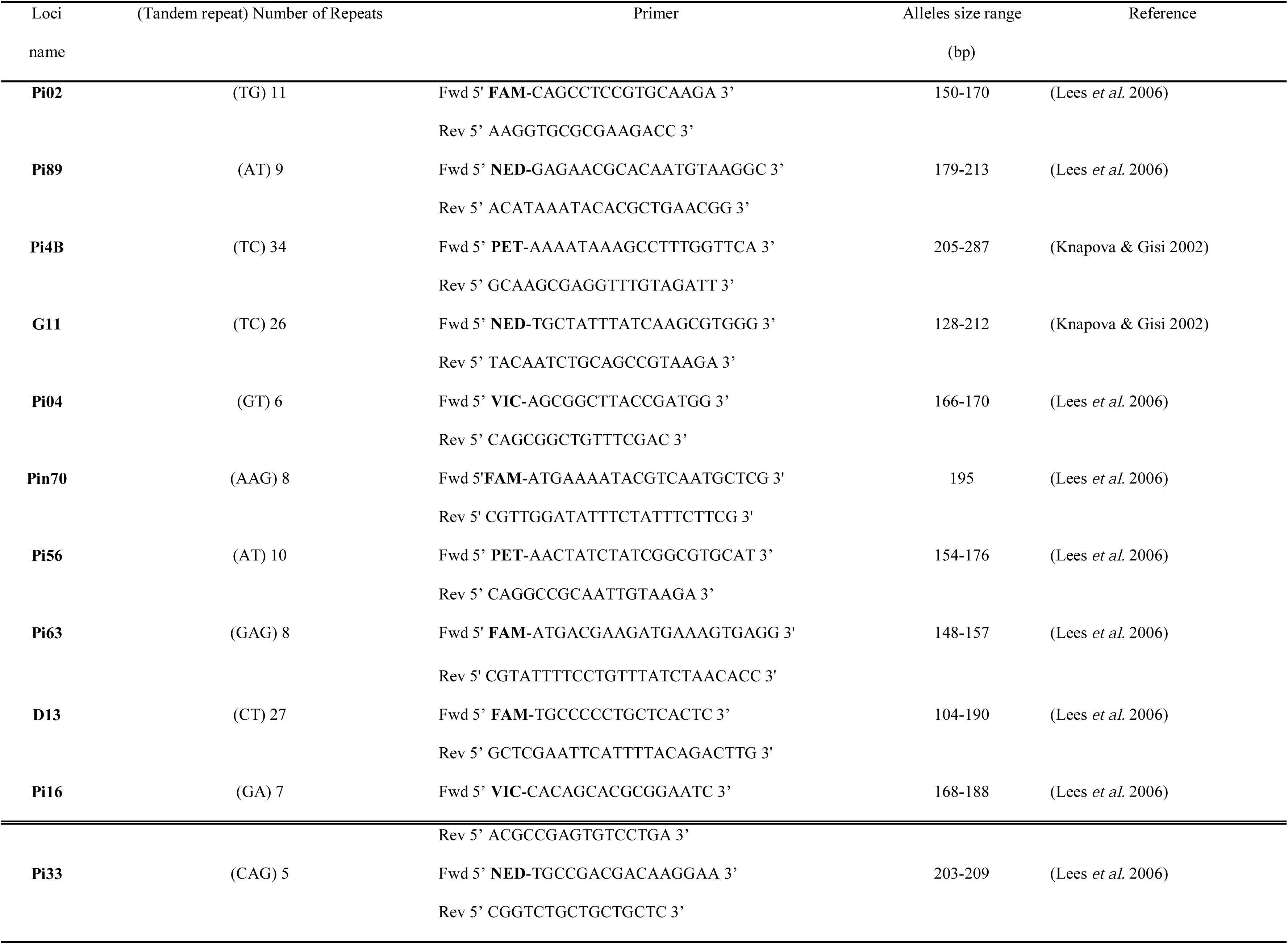
List of SSR markers and allele size range.

**Table S3.** Shapiro-Wilk tests to assess normality for morphological traits (P = 0.05).

|  | Shapiro-Wilk Test |  |
| --- | --- | --- |
|  | W statistics | p-value |
| **Mycelia radial growth (cm<sup>2</sup>)** | 0.6946 | $1.0 \times 10^{-15}$ |
| **Hyphal width (μm)** | 0.9685 | $1.0 \times 10^{-15}$ |
| **Length of sporangia (μm)** | 0.9937 | $1.1 \times 10^{-5}$ |
| **Width of sporangia (μm)** | 0.8448 | $1.0 \times 10^{-15}$ |
| **Length:width ratio (μm)** | 0.9684 | $1.0 \times 10^{-15}$ |
| **Area of sporangia (μm<sup>2</sup>)** | 0.9530 | $1.0 \times 10^{-15}$ |

**Table S4** STRUCTURE Δ*K* values for all 100 resampled datasets for ploidy = 2.

**Table S5** STRUCTURE Δ*K* values for all 100 resampled datasets for ploidy = 3.

**Table S6.** Mean, minimum, and maximum percentage variance explained by each Principal Component (PC) across the 100 resampled datasets for ploidy = 2 and ploidy = 3.

| Ploidy | PC | Mean variance explained | Minimum variance explained | Maximum variance explained |
| --- | --- | --- | --- | --- |
| 2 | 1 | 11.93% | 11.70% | 12.05% |
| 2 | 2 | 4.69% | 4.54% | 4.89% |
| 2 | 3 | 4.11% | 4.05% | 4.14% |
| 2 | 4 | 3.70% | 3.63% | 3.90% |
| 2 | 5 | 3.65% | 3.60% | 3.68% |
| 3 | 1 | 12.94% | 12.65% | 13.38% |
| 3 | 2 | 4.99% | 4.87% | 5.13% |
| 3 | 3 | 4.03% | 3.92% | 4.30% |
| 3 | 4 | 3.85% | 3.75% | 4.05% |
| 3 | 5 | 3.75% | 3.67% | 3.87% |

**Table S7.** Effect of temperature (4, 18, and 25°C) and culture media (V8 juice agar (V8), Potato Dextrose Agar (PDA), Corn Meal Agar (CMA), and Tree Tomato Agar (TTA)) on average mycelial radial growth for five isolates of *P. betacei,* seven isolates of *Phytophthora infesta*ns, and three isolates of *P. andina* (EC-2 and EC3 clonal lineages). Values were assessed after 15 days of incubation.

|  | *P. betacei* | *P. infestans* | *P. andina* (EC-2) | *P. andina* (EC-3) |
| --- | --- | --- | --- | --- |
| **Growth temperature (°C)** |  |  |  |  |
| **Minimum and maximum temperature (°C)** | 18 °C<sup>1</sup> | 18 °C to 25 °C | 18 °C to 25 °C | 18 °C<sup>1</sup> |
| **Optimum temperature (°C)** | 18 °C | 18 °C | 18 °C | 18 °C |
| **Average colony growth at 4 °C (mm)** |  |  |  |  |
| **V8 juice agar** | No growth | No growth | No growth | No growth |
| **Potato Dextrose Agar (PDA)** | No growth | No growth | No growth | No growth |
| **Corn Meal Agar (CMA)** | No growth | No growth | No growth | No growth |
| **Tree Tomato Agar (TTA)** | No growth | No growth | No growth | No growth |
| **Average colony growth at 18 °C (mm)** |  |  |  |  |
| **V8 juice agar** | 37.77 (±9.62) | 24.08 (±19.60) | 22.26(±16.81) | 28.15 (±19.08) |
| **Potato Dextrose Agar (PDA)** | 0.27 (±0.51) | 37.87 (±8.66) | 38.72 (±5.01) | 38.78 (±0.47) |
| **Corn Meal Agar (CMA)** | 22.88 (±12.65) | 21.05 (±9.49) | 30.27 (±5.24) | 24.00 (±7.48) |
| **Tree Tomato Agar (TTA)** | 34.63 (±6.15) | 14.11 (±9.66) | 7.26(3.84) | 15.21 (±1.51) |
| **Average colony growth at 25 °C (mm)** |  |  |  |  |
| **V8 juice agar** | No growth | 5.33(±17.95) | 2.18(±3.81) | No growth |
| **Potato Dextrose Agar (PDA)** | No growth | 6.97 (±10.55) | 10.09(±9.72) | No growth |
| **Corn Meal Agar (CMA)** | No growth | 0.48 (±1.87) | No growth | No growth |
| **Tree Tomato Agar (TTA)** | No growth | 5.58(±5.87) | 0.39(±0.10) | No growth |

<sup>1</sup> For *P. betacei* isolates, no mycelial growth was observed at either 4 °C or 25 °C.

**Table S8.** Effect of temperature on average mycelial radial growth (cm^2^) for *Phytophthora betacei*, *Phytophthora infestans*, and Phytophthora andina isolates on V8 juice agar (V8), Potato Dextrose Agar (PDA), Corn Meal Agar (CMA), and Tree Tomato Agar (TTA).

| Isolates | Average mycelia radial growth (cm<sup>2</sup>) (standard deviation) |  |  |  |  |  |  |  |  |  |  |  |
| --- | --- | --- | --- | --- | --- | --- | --- | --- | --- | --- | --- | --- |
|  | 4 °C |  |  |  | 18 °C |  |  |  | 25 °C |  |  |  |
|  | V8 agar | PDA | CMA | TTA | V8 agar | PDA | CMA | TTA | V8 agar | PDA | CMA | TTA |
| *P. betacei* | 0.00 | 0.00 | 0.00 | 0.00 | 37.77 (±9.62) | 0.27 (±0.51) | 22.88 (±12.65) | 33.02 (±6.15) | 0.00 | 0.00 | 0.00 | 0.00 |
| *P. andina* | 0.00 | 0.00 | 0.00 | 0.00 | 24.22 (±16.96) | 38.76 (±4.00) | 28.18 (±6.50) | 7.97(±5.54) | 1.72 (±3.15) | 6.54 (±9.34) | 0.05 (±0.12) | 0.00 |
| *P. infestans* | 0.00 | 0.00 | 0.00 | 0.00 | 24.08 (±19.60) | 37.87 (±8.66) | 21.05 (±9.49) | 13.25(±9.66) | 5.33(±17.95) | 6.97 (±10.55) | 0.48 (±1.87) | 0.00 |

**Table S9.**
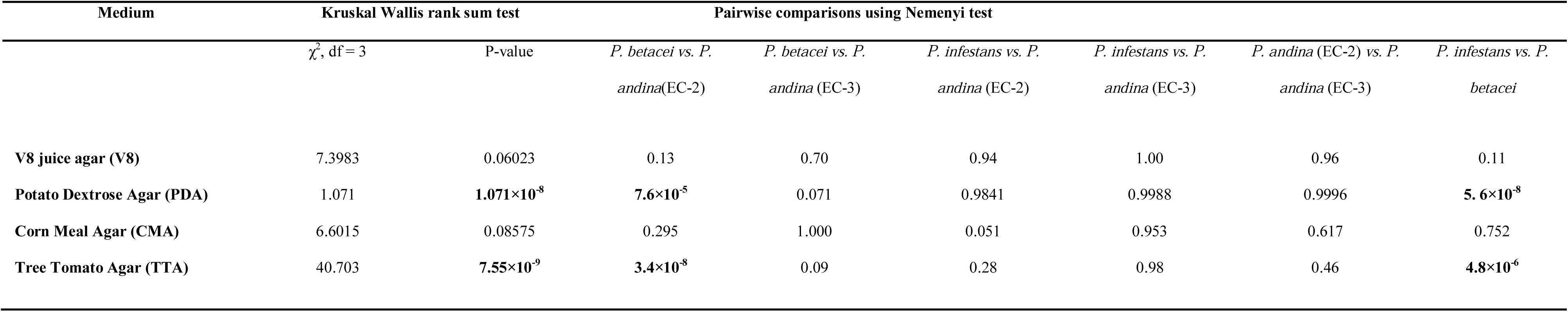
Mycelial radial growth differs among species within the *Phytophthora infestans* complex. Kruskal-Wallis rank sum test followed by pairwise comparisons using the Nemenyi test for mycelial radial growth at 18°C on V8 juice agar (V8), Potato Dextrose Agar (PDA), Corn Meal Agar (CMA), and Tree Tomato Agar (TTA). Significant P-values (P < 0.05) are shown in bold.

| Medium | Kruskal Wallis rank sum test |  | Pairwise comparisons using Nemenyi test |  |  |  |  |  |
| --- | --- | --- | --- | --- | --- | --- | --- | --- |
| | $\chi^2$, df = 3 | P-value | *P. betacei* vs. *P. andina*(EC-2) | *P. betacei* vs. *P. andina* (EC-3) | *P. infestans* vs. *P. andina* (EC-2) | *P. infestans* vs. *P. andina* (EC-3) | *P. andina* (EC-2) vs. *P. andina* (EC-3) | *P. infestans* vs. *P. betacei* |
| V8 juice agar (V8) | 7.3983 | 0.06023 | 0.13 | 0.70 | 0.94 | 1.00 | 0.96 | 0.11 |
| Potato Dextrose Agar (PDA) | 1.071 | **1.071×10<sup>-8</sup>** | **7.6×10<sup>-5</sup>** | 0.071 | 0.9841 | 0.9988 | 0.9996 | **5.6×10<sup>-8</sup>** |
| Corn Meal Agar (CMA) | 6.6015 | 0.08575 | 0.295 | 1.000 | 0.051 | 0.953 | 0.617 | 0.752 |
| Tree Tomato Agar (TTA) | 40.703 | **7.55×10<sup>-9</sup>** | **3.4×10<sup>-8</sup>** | 0.09 | 0.28 | 0.98 | 0.46 | **4.8×10<sup>-6</sup>** |

**Table S10.** Comparisons of all quantitative morphological traits (length, width, length:width, and area) of sporangia between P. betacei, P. infestans, and P. andina (Clonal lineage EC-2 and EC-3) isolates growing at 18°C on V8 juice agar (V8), Potato Dextrose Agar (PDA), Corn Meal Agar (CMA), and Tree Tomato Agar (TTA). Means of all data are followed by the Kruskal-Wallis rank sum test results and a pairwise comparisons using Nemenyi test. Statistically significant P-values (P < 0.01) are shown in bold.

|  |  | Mean |  |  |  |
| --- | --- | --- | --- | --- | --- |
|  |  | *P. betacei* | *P. andina* (EC-2) | *P. andina* (EC-3) | *P. infestans* |
| **Length sporangia**<br>($\mu\text{m}$) | V8 | 36.32($\pm 5.99$) | 34.24 ($\pm 5.55$) | 34.34(6.84) | 33.30 ($\pm 5.96$) |
| | PDA | No growth | 36.38 ($\pm 3.76$) | 36.10($\pm 5.77$) | 31.33 ($\pm 5.00$) |
| | CMA | 37.61 ($\pm 5.42$) | 36.16 ($\pm 4.81$) | 35.91($\pm 3.27$) | 32.02 ($\pm 4.80$) |
| | TTA | 39.34 ($\pm 4.77$) | 37.89 ($\pm 4.33$) | 36.30($\pm 6.06$) | 31.55 ($\pm 4.98$) |
| **Width of sporangia**<br>($\mu\text{m}$) | V8 | 17.26($\pm 2.92$) | 18.50 ($\pm 2.98$) | 18.66($\pm 3.88$) | 18.53($\pm 3.34$) |
| | PDA | No growth | 18.50 ($\pm 1.855$) | 18.28($\pm 3.03$) | 20.07($\pm 3.37$) |
| | CMA | 16.93($\pm 2.34$) | 17.55 ($\pm 2.27$) | 17.50($\pm 1.67$) | 19.18($\pm 2.88$) |
| | TTA | 15.75($\pm 5.72$) | 18.13 ($\pm 2.67$) | 17.39($\pm 2.75$) | 20.67($\pm 3.42$) |
|  | V8 | 2.13(±0.41) | 1.85 (±0.044) | 1.85(0.049) | 1.81(±0.20) |
| **Length/width ratio** | PDA | No growth | 1.96 (±0.05) | 1.97(±0.05) | 1.56(±0.10) |
| **(µm)** | CMA | 2.22(±0.20) | 2.05 (± 0.021) | 2.05(±0.024) | 1.66(±0.01) |
|  | TTA | 2.56(±0.19) | 2.11 (±0.18) | 2.10(±0.19) | 1.53(±0.08) |
|  | V8 | 436.28(±119.38) | 511.08 (±23.97) | 510.01(±24.36) | 550.03(±141.57) |
| **Mean of sporangia** | PDA | No growth | 467.94 (±13.90) | 464.62(±13.54) | 665.98(±112.79) |
| **area (µm<sup>2</sup>)** | CMA | 386.12(±33.11) | 442.14(±9.03) | 444.47 (±10.85) | 576.82(±9.10) |
|  | TTA | 311.46(±39.53) | 420.95 (±63.52) | 424.36(±66.62) | 692.79(±107.51) |

| Kruskal Wallis rank sum test |  |  |  | Pairwise comparisons using Nemenyi test |  |  |  |  |  |
| --- | --- | --- | --- | --- | --- | --- | --- | --- | --- |
| | | $\chi^2$,<br>df = 3 | P | *P. betacei* vs. *P.*<br>*andina*(EC-2) | *P. betacei* vs. *P.*<br>*andina*(EC-3) | *P. infestans* vs. *P.*<br>*andina*(EC-2) | *P. infestans* vs. *P.*<br>*andina*(EC-3) | *P. betacei* vs. *P.*<br>*infestans* | *P. andina* (EC-2) vs.<br>*P. andina* (EC-3) |
| Length sporangia<br>($\mu\text{m}$) | V8 | 19.134 | 0.0002565 | 0.09 | 0.32 | 0.88 | 0.92 | **0.00013** | 0.99 |
| | PDA | 60.52 | $7.19 \times 10^{-14}$ | NA | NA | **$1.7 \times 10^{-11}$** | **$7 \times 10^{-11}$** | NA | 0.92 |
| | CMA | 41.999 | $4.014 \times 10^{-9}$ | 0.68 | 0.6626 | **0.00061** | **0.01405** | **$6.0 \times 10^{-10}$** | 0.99 |
| | TTA | 144.86 | $2.2 \times 10^{-16}$ | 0.39 | **0.03656** | **$8.2 \times 10^{-12}$** | **0.00083** | **$< 1 \times 10^{-15}$** | 0.53 |
| Width of sporangia<br>($\mu\text{m}$) | V8 | 16.209 | 0.001027 | 0.367 | 0.2037 | 0.9999 | 1.0000 | **0.0017** | 0.9999 |
|  | PDA | 11.836 | 0.002691 | NA | NA | **0.0093** | **0.0622** | NA | 0.9988 |
| | CMA | 25.211 | $1.39 \times 10^{-5}$ | 0.451 | 0.596 | **0.045** | 0.190 | **$3.4 \times 10^{-6}$** | 1.000 |
| | TTA | 174.53 | $< 2.2 \times 10^{-16}$ | **$8.1 \times 10^{-8}$** | **0.00794** | **0.00024** | **0.00018** | **$< 2 \times 10^{-16}$** | 0.76033 |
| Length/width ratio<br>($\mu\text{m}$) | V8 | 11.584 | 0.008951 | 0.1876 | 0.4597 | 0.9886 | 0.9913 | **0.0069** | 1.0000 |
| | PDA | 188.47 | $< 1 \times 10^{-15}$ | NA | NA | **$< 2.0 \times 10^{-16}$** | **$3.8 \times 10^{-14}$** | NA | 0.95 |
| | CMA | 210.76 | $< 2.2 \times 10^{-15}$ | **$7.2 \times 10^{-13}$** | **$4.1 \times 10^{-9}$** | **0.0014** | **0.0246** | **$< 1 \times 10^{-15}$** | 0.99 |
| | TTA | 350.4 | $< 2 \times 10^{-16}$ | **$1.5 \times 10^{-10}$** | **$2.5 \times 10^{-6}$** | **$2.8 \times 10^{-12}$** | **$4.3 \times 10^{-7}$** | **$< 1 \times 10^{-15}$** | 1 |
| Mean of sporangia<br>area ($\mu\text{m}^2$) | V8 | 10.863 | 0.01249 | 0.19 | 0.51 | 0.99 | 0.99 | **0.01** | 1.00 |
| | PDA | 188.47 | $< 2 \times 10^{-16}$ | NA | NA | **$< 2.0 \times 10^{-16}$** | **$< 3.8 \times 10^{-14}$** | NA | 0.95 |
| | CMA | 210.75 | $< 2 \times 10^{-16}$ | **$7.2 \times 10^{-13}$** | **$4.2 \times 10^{-9}$** | **0.0014** | **0.02** | **$< 1 \times 10^{-15}$** | 0.99 |
| | TTA | 350.4 | $< 2 \times 10^{-16}$ | **$8.6 \times 10^{-14}$** | **$1.5 \times 10^{-10}$** | **$2.8 \times 10^{-12}$** | **$4.3 \times 10^{-7}$** | **$< 2 \times 10^{-16}$** | 1 |

**Table S11.**
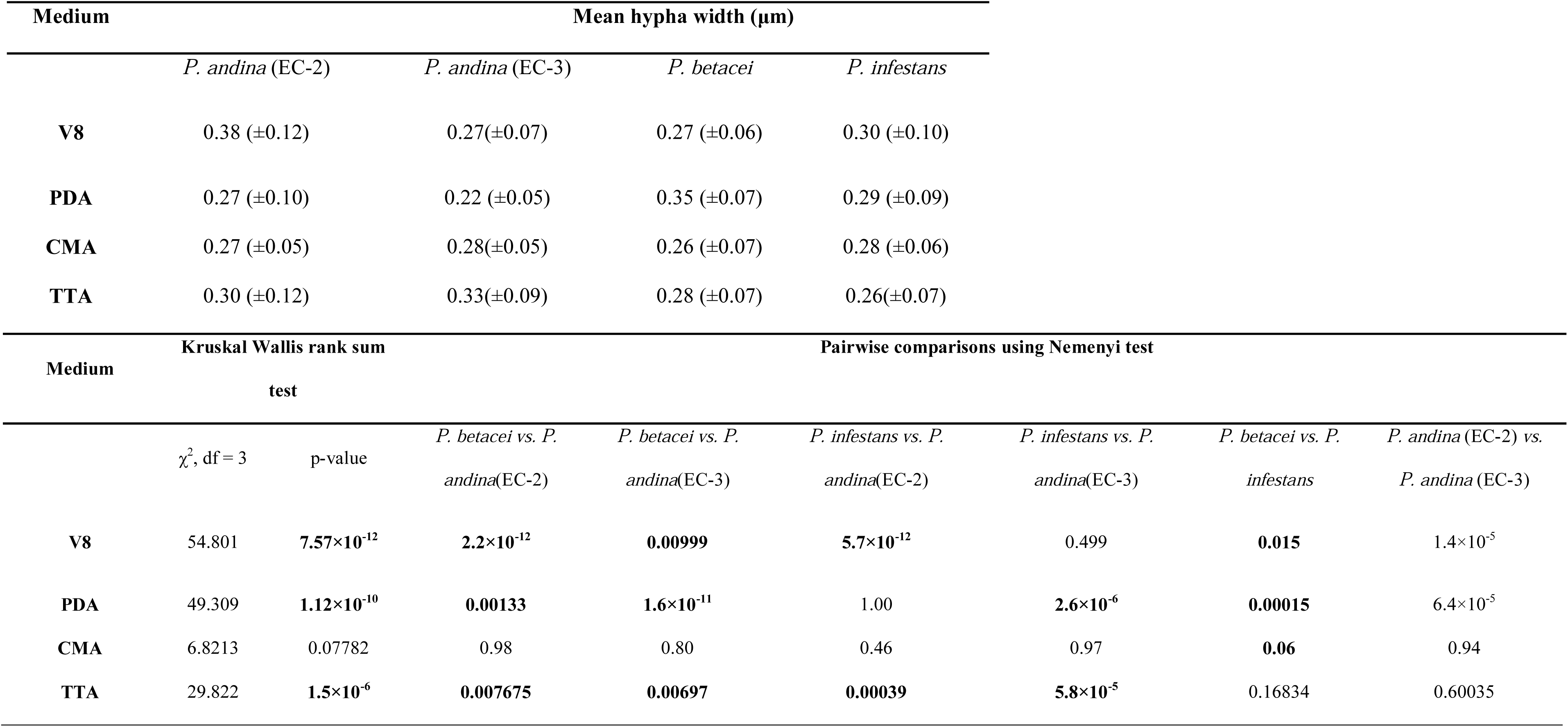
**Differences in hyphal width between *P. betacei, P. andina* (Clonal lineage EC-2 and EC-3), and *P. infestans* isolates growing at 18° C on four different media.** Media assessed were V8 juice agar (V8), Potato Dextrose Agar (PDA), Corn Meal Agar (CMA), and Tree Tomato Agar (TTA). Means of all data are followed by Kruskall-Wallis rank sum test. For all pairwise comparisons Nemenyi test or Tukey test is shown. Statistically significant P-values (P < 0.05) are shown in bold.

## SUPPLEMENTARY FIGURES

**Figure S1. Geographic distribution of sample collection sites.** *P. betacei* strains isolated from tree tomato were collected from several sampling locations, which are indicated with yellow circles. A complete list of each sampling site along with the geographic coordinates is shown in Table S1.

**Figure S2. Disease symptoms on tree tomato (*Solanum betaceum*) associated with *P. betacei*.** (**A**) Late blight infection and typical symptoms on leaf caused by *P. betacei* in the field. (**B**) Damage and necrotic lesion caused by *P. betacei* on stems of tree tomato plants. (**C**) Typical symptoms on leaf caused by *P. infestans* in cultivated potato fields. (**D**) Tree tomato plants without infections. (**E**) Devastating effect of *P. betacei* on tree tomato fields five days after symptoms first appeared. (**F**) Devastating effect of *P. infestans* on cultivated potato fields two days after symptoms first appeared.

**Figure S3. Principal component analysis (PCA) for *P. betacei, P. infestans* and *P. andina* using resampled SSR datasets.** Principal components (PC) 1 and 2 are shown and the percentage of variance explained by each eigenvalue is shown within parentheses on each axis. Individuals of *P. infestans* are shown in blue, *P. andina* in orange and *P. betacei* in green. (**A**) PCA results for all 100 diploid resampled datasets, and (**B**) PCA results for all 100 triploid resampled datasets. The principal component values were not greatly affected by uncertainty in allele frequencies at bi-allelic triploid loci. In all cases PC1 differentiates between *P. infestans sensu stricto* and *P. betacei*. Each point represents a resampled dataset.

**Figure S4. Principal component analysis (PCA) using 12 microsatellite loci.** *P. infestans* isolates are shown in blue, *P. andina* in orange and *P. betacei* in green. Principal component results are plotted for all 100 resampled datasets for each ploidy. Ploidy = 2 and ploidy = 3. Mean of variance explained Ploidy = 2 and ploidy = 3 (Table S12).

**Figure S5. Genetic structure among *P. infestans, P. betacei* and *P. andina* samples using Genotyping-by-sequencing data.** (A) Classification of 48 *Phytophthora* samples into tree different populations according to the optimal population number (Δ*K* = 3; Evanno’s method). The distribution of the individuals in different populations is indicated by the color code (purple= *P. infestans,* blue=P. *betacei* and yellow *P. andina* (EC-2)) (B) The estimated posterior probability, log likelihood of the data, for a given *K* (*LnP*(D)) and ad hoc quantity *ΔK* computed for GBS data for 6 populations (K = 1 to 6).

**Figure S6. STRUCTURE results for one resampled dataset with ploidy = 2 and *K* = 1 through *K* = 8.** Δ*K* values for each of the 100 resampled diploid datasets are shown in Table S13.

**Figure S7. STRUCTURE results for one resampled dataset with ploidy = 3 and *K* = 1 through *K* = 8.** Δ*K* values for each of the 100 resampled triploid datasets are shown in Table S14.

**Figure S8. *STRUCTURE* analyses are robust to uncertainty in allele frequencies at bi-allelic triploid sites in *P. infestans* tested.** Boxplots showing the variation in population assignment for *K* = 2 for each isolate across all 100 resampled datasets. Each individual is represented by a vertical column containing two boxes showing the respective variation in assignment to the two detected populations. Boxes represent the 25 to −75 quantiles of the population assignments with whiskers extending to minimum and maximum values. The results show little overall variance in population assignment. (**A**) Ploidy = 2. (**B**) Ploidy = 3.

**Figure S9. Comparative colony morphology, mycelial radial growth and sporangia morphology of *Phytophthora andina, Phytophthora betacei* and *Phytophthora infestans*.**
Differences on colony morphology and mycelial radial growth on all media tested: V8 juice agar (**A**), Potato Dextrose Agar (**B**), Corn Meal Agar (**C**), and Tree Tomato Agar (**D**) after 7 days (left) and 15 days (right) of incubation at 18°C.

**Figure S10. Mycelial radial growth among the *Phytophthora betacei* tested at 4, 18 and 25°C.** Effect of temperature and media on mycelial radial growth on V8 juice agar (V8), Potato Dextrose Agar (PDA), Corn Meal Agar (CMA), and Tree Tomato Agar (TTA) of *P. betacei, P. infestans,* and *P. andina* isolates 15 days after incubation at three different temperatures (4, 18, and 25°C).

**Figure S11. Colony growth and sporangia morphology of *Phytophthora betacei* after 15-days of incubation at optimum growth temperature (18°C).** Colony growth of *P. betacei* was evaluated on on the V8 juice agar (V8), Potato Dextrose Agar (PDA), Corn Meal Agar (CMA), and Tree Tomato Agar (TTA) after 15 days of incubation at optimum growth temperature (18°C) (**A-D**). Sporangia borne terminally to the sporangiophore, caducous, ovoid and semi-papillate. Pictures of sporangia on V8 juice agar (V8), Potato Dextrose Agar (PDA), Corn Meal Agar (CMA) and Tree Tomato Agar (TTA) (**E-H**). Scaled bar = 50 μm.

**Figure S12. Effect of media on hyphal width (μm) among the *Phytophthora* species.** Hyphal width was tested for all isolates on V8 juice agar (V8), Potato Dextrose Agar (PDA), Corn Meal Agar (CMA), and Tree Tomato Agar (TTA) after 15-days of incubation at optimum growth temperature (18°C).

